# Glypican-5 delineates spatially distinct astrocyte and neuronal populations in the human hippocampus and is selectively remodelled in Alzheimer’s disease

**DOI:** 10.1101/2025.08.19.671074

**Authors:** Félicia Jeannelle, Mónica Miranda de la Maza, Sophie Schreiner, Gaël Paul Hammer, Dominique Mirault, Naguib Mechawar, Brainbank Neuro-CEB Neuropathology Network, Netherlands Brain Bank, Michel Mittelbronn, David S. Bouvier

## Abstract

Regional heterogeneity of astrocytes and neurons is increasingly recognised as a determinant of selective vulnerability in neurodegeneration, yet the molecular signatures underlying this specificity remain poorly defined. Glypican-5 (GPC5), a heparan sulfate proteoglycan expressed mainly by astrocytes, contributes to synaptic organisation and circuit stability, whose disruption may undermine astrocyte-neuron crosstalk and contribute to selective neuronal loss in neurodegenerative diseases. Using multiplex chromogenic immunohistochemistry, in situ hybridization and digital pathology, we mapped GPC5 expression across the hippocampus and parahippocampal cortex in post-mortem tissue from non-demented control (CTL), Alzheimer’s disease (AD) and Parkinson’s disease with dementia (PDD) cases.

In CTL brains, GPC5 labelled spatially restricted populations of astrocytes and pyramidal neurons organized according to hippocampal subfield and laminar architecture. GPC5-positive astrocytes co-expressed canonical markers but represented a more restricted population while GPC5 protein was enriched at synapse-rich regions of the outer molecular layer of the dentate gyrus. In AD and PDD, regional distribution patterns of GPC5 were distinct from canonical astrocyte markers including GFAP, AQP4 and ALDH1L1. In PDD, GPC5 distribution was largely preserved. In AD, GPC5 underwent selective redistribution with a significant loss of the staining in the dentate gyrus, and an accumulation on amyloid plaques, putatively secreted by plaque-associated astrocytes, and on neurofibrillary tangles, likely of neuronal and astrocytic origin

These findings reveal disease- and region-specific remodelling of a spatially organised astrocyte-neuron system and establish GPC5 as a molecularly distinct responder to AD pathology.

## INTRODUCTION

Selective regional vulnerability is a defining feature of neurodegenerative diseases (NDDs). In Alzheimer’s or Parkinson’s disease, specific neuronal populations are affected early on, including noradrenergic neurons of the locus coeruleus (LC), dopaminergic neurons of the substantia nigra pars compacta (SNc), as well as some pyramidal neurons of the entorhinal cortex and Cornu Ammonis 1 (CA1) region of the hippocampus^1^. The mechanisms underlying this selective vulnerability remain incompletely understood and are often investigated from a neuronal perspective. However, there is accumulating evidences that astrocytic molecular and functional heterogeneity might also contribute to regional disease progression. Accounting for 20–40% of all brain cells, astrocytes support neuronal function through an extensive repertoire of tasks, including the regulation of synaptic formation and transmission, neurotransmitter and ion homeostasis^2–4–3^, metabolic coupling, blood–brain barrier integrity, cerebral blood flow^5,6^, and glymphatic clearance^7^. Their highly ramified morphology enables a tiled organization of the parenchyma, allowing individual astrocytes to monitor and modulate thousands of synapses^8^ while engaging the vasculature through specialized endfeet. Far from being a homogeneous population, astrocytes display marked molecular, morphological, and functional diversity across brain regions, particularly in the human brain^9,10^. This heterogeneity has important consequences for neuronal circuit organization and function. During development, astrocyte molecular identity is shaped by the local neuronal environment^11^, and in turn astrocytes play essential roles in synapse formation, maturation, and maintenance^12,13^. Astrocytes also undergo profound molecular and functional remodelling in response to injury, inflammation, or protein aggregation^14–17^. While some responses are neuroprotective and support tissue repair, sustained or maladaptive reactivity can promote neurotoxic states and accelerate neurodegenerative cascades^17^. Recent single-cell transcriptomic studies have further highlighted the complexity of astrocyte responses in neurodegeneration, revealing multiple disease-associated and homeostatic astrocyte states. Among these, disease-associated astrocytes (DAAs) emerge early in AD and are characterized by increased expression of reactive markers, including GFAP^18,19^. In contrast, astrocyte populations displaying lower GFAP expression exhibit distinct transcriptional signatures that remain comparatively less explored. One such marker is glypican-5 (GPC5), a glycosylphosphatidylinositol-anchored heparan sulfate proteoglycan that has emerged as a candidate regulator of astrocyte–neuron interactions^20^. GPC5 is expressed in the mammalian brain throughout development and adulthood^21^ and is particularly enriched in astrocytes where it has been implicated in synapse maturation and maintenance^22^. Recent evidence further suggests a role for GPC5 in neurodegeneration. In the APP/PS1 mouse model of AD, GPC5 expression is reduced, whereas astrocyte-specific overexpression rescues early synaptic dysfunction and improves cognition, supporting a potential neuroprotective function^23^. Dysregulation of GPC5 has also been reported in various NDDs, including fronto-temporal dementia^24^, multiple sclerosis^25^ and AD^26,27^, suggesting broader relevance to neurodegeneration and neuroinflammation. Despite these observations, the anatomical distribution of GPC5-positive astrocytes in the human brain and their relationship to neurodegenerative pathology remain poorly understood. It remains unclear whether GPC5 defines a distinct astrocyte subtype, how it relates to established astrocytic and disease-associated markers, and whether its expression is altered in AD and Parkinson’s disease with dementia (PDD). Addressing these questions is essential to determine whether GPC5 identifies a specialized astrocyte population potentially involved in regional vulnerability and resilience to neurodegeneration. In this study, we systematically map *GPC5* mRNA and protein expression across cell types and disease conditions in the human hippocampus and parahippocampal cortex, brain regions critically affected during early memory decline. We further assess its relationship with established astrocytic markers as well as neuronal populations to refine our understanding of its role in both physiological and pathological conditions.

## MATERIAL AND METHODS

### Human brain samples

Post-mortem formalin-fixed paraffin embedded (FFPE) hippocampal and cortical tissues were obtained from The Netherlands Brain Bank (NBB, Amsterdam, The Netherlands), the Douglas-Bell Canada Brain Bank (Douglas Mental Health University Institute, Montreal, QC, Canada), and the GIE-Neuro-CEB biobank (Groupe Hospitalier Pitié-Salpêtrière, Paris, France). Research involving these samples was conducted with approval from the respective ethic committees of these institutions, as well as the University of Luxembourg (ERP 16-037 and 21-009).

Neuropathological assessments were carried out by specialized neuropathologists at the brain banks, following standardized criteria including Braak^28,29^, ABC^30^ and McKeith^31^ staging, based on the presence of Aβ plaques, neurofibrillary tangles (NFTs), and α-synuclein pathology.

FFPE hippocampal and cortical samples include age-matched non-demented CTLs (n=15), AD (n= 26) and PDD patients (n=10). Age at death, post-mortem delay (PMD), sex and staging of the human subjects are listed in Supplementary Table 1.

### Immunohistochemistry (IHC)

FFPE hippocampal and cortical tissue blocs were sectioned at 5 µm thickness using a standard microtome and mounted on Dako FLEX IHC-coated slides (Cat# K8020, Agilent). The sections were then dried at 60°C for at least one hour before undergoing automated staining on a Dako Omnis Immunostainer (Agilent). Antigen retrieval was performed using heat-induced epitope retrieval (HIER) with high- or low-pH buffer solutions (EnVisionTMFLEX Target Retrieval Solution, Cat# K8004 and Cat# K8005, respectively) for 30 min at 97°C.

Primary antibodies (anti-ALDH1L1 Cat# HPA050139 and anti-AQP4 Cat# AMAb90537) were prepared in EnVisionTMFLEX antibody diluent (Cat# K8006), with specific concentrations provided in Supplementary Table 2. The sections were incubated with primary antibodies for one hour at room temperature (RT), followed by visualisation using a horseradish peroxidase (HRP)/3,3′-diaminobenzidine (DAB) detection system (EnVisionTMFLEX Detection Kit, Cat# K8000). Counterstaining was performed using haematoxylin (Cat# GC808), after which the slides were dehydrated through ethanol washes and mounted.

### Single and Multiplex chromogenic IHC (cIHC)

For experiments conducted using the Ventana Discovery Ultra automated IHC and *in situ* hybridisation (ISH) platform (Roche Ventana Medical Systems, Tucson, AZ, USA), 5 µm sections from FFPE tissue blocs were mounted onto hydrophilic adhesion slides (Matsunami TOMO®, Cat# TOM-1190). Slides were dried as previously described. All reagents were provided within the system as per Roche Diagnostics’ recommendations, and washing steps were carried out using Reaction Buffer (Cat# 950-300). Antibody concentrations used in the assays are detailed in Supplementary Table 2.

For both single and multiplex IHC, counterstaining was performed using haematoxylin II (Cat# 790-2208) and bluing reagent (Cat# 760-2039), each incubated for 4 min. Following staining, all slides underwent washing, dehydration through a graded alcohol series, and were coverslipped.

For single-plex cIHC, sections were first deparaffinised at 69°C in EZ Prep solution (Cat# 950-102) for 8 min, repeated across three cycles. HIER was carried out at 95°C for 40 min using Cell Conditioning 1 (CC1, Cat# 950-224). To block endogenous peroxidase and non-specific protein interactions, sections were incubated with Chromomap (CM) inhibitor at 37°C for 8 min. Primary antibodies (anti-ALDH1L1 Cat# HPA050139, anti-ALDH7A1 Cat# HPA023296, anti-AQP4 Cat# AMAb90537, anti-GFAP Cat# 760-4345, anti-GPC5 Cat# HPA040152, anti-VGAT Cat# HPA058859 and anti-VGLUT1 Cat# HPA063679) were automatic and manually applied after dilution in EnVisionTMFLEX buffer, respectively. Incubation with primary antibodies lasted 60 min, followed by detection using OmniMap anti-Rb (Cat# 760-4311) or OmniMap anti-Ms (Cat# 760–4310) for 16 min. This was followed by sequential incubations of H_₂_O_₂_ CM (4 min), DAB CM (8 min), and Copper CM (4 min), all from the Discovery CM DAB kit (Cat# 760-159). All multiplex protocols (two- to three-plex) shared a common initial sequence, including deparaffinization, HIER using CC1, and blocking with the Discovery inhibitor, unless otherwise specified.

In the two-plex combinations (GPC5/GFAP, GPC5/ALDH1L1, GPC5/NeuN, ALDH1L1/GFAP, AQP4/GFAP, GPC5/AQP4, ALDH7A1/GPC5, ALDH7A1/GFAP, ALDH1L1/ALDH7A1, GPC5/AT8 and GPC5/ps396) following deparaffinization and HIER, one drop of Discovery inhibitor (Cat# 760-4840) was applied for 8 min. Subsequently, primary antibodies including anti-GPC5, anti-ALDH1L1, anti-ALDH7A1 and anti-AQP4 were incubated for 60 min. These were developed using OmniMap anti-Rb or OmniMap anti-Ms HRP for 16 min, followed by a chromogenic reaction using Discovery Purple and H_2_O_2_ Purple from the Discovery Purple Kit (Cat# 760-229), with 4 min and 32 min of incubation, respectively. Between each staining cycle, heat-mediated antibody denaturation (100°C for 24 min) was performed using Cell Conditioning 2 (CC2) reagent (Cat# 950-223) to dissociate primary antibody-HRP complexes and prevent cross-reactivity with residual HRP. Next, anti-GFAP, anti-ALDH1L1, anti-ALDH7A1, anti-NeuN (Cat# MAB377), anti-phospho tau AT8 (Cat# MN1020), anti-phospho tau pS396 (Cat# 44-752G), anti-GPC5, and anti-AQP4 were applied for 48 min (anti-GFAP) or 60 min (all others). Detection involved signal amplification with OmniMap anti-Rb or anti-Ms HRP for 16 min, followed by chromogenic development with either Discovery Teal HRP chromophore (Cat# 760-247) or Discovery Green HRP chromophore (Cat# 760-271), involving a 4-min incubation with Teal/Green HRP Substrate, 32 min with Teal/Green HRP H_2_O_2_, and 16 min with Teal/Green HRP Activator.

The three-plex protocols (4G8/GPC5/GFAP; pSYN/GPC5/GFAP) were carried out as follows: anti-4G8 (Cat# 800712) or anti-pSYN (Cat# MABN826) were incubated for 1 hour and 32 min, amplified with OmniMap anti-Ms HRP for 16 min, and revealed using Discovery CM DAB according to the single-plex IHC method. After denaturation, anti-GPC5 was applied for 60 min, followed by OmniMap anti-Rb HRP for 16 min and chromogenic detection with Discovery Purple. Finally, anti-GFAP was incubated for 48 min, followed by signal amplification using OmniMap anti-Rb HRP for 16 min and development using either Discovery Teal HRP or Discovery Green HRP.

### Multiplex chromogenic In situ hybridization (ISH) and IHC

RNAscope multiplex chromogenic ISH and IHC was performed using the Ventana Discovery Ultra automated staining platform (Ventana Medical Systems, Tucson, USA) and the mRNA Universal Procedure (v8.00) workflow. The protease-free protocol was applied to preserve protein epitopes for downstream IHC and it was achieved by using RNAscope VS Pro Reagents (ACD/Bio-Techne, Cat# 322035). All reagents were provided within the system as per Roche Diagnostics’ and ACD/Bio-Techne’s recommendations, and washing steps were carried out using Reaction Buffer (Roche, Cat# 950-300).

5 µm FFPE tissue sections were mounted onto hydrophilic adhesion slides (Matsunami TOMO®, Cat# TOM-1190). Slides were dried as previously described. Slides were deparaffinized at 60°C for 32 min and rehydrated at 58°C using EZ Prep solution (Cat# 950-102) for 8 min, repeated for three cycles. Dewaxing was then carried out using RNAscope VS Dewax mRNA Dewax (ACD/Bio-Techne, RNAScope VS Dewax, Cat# 323222). HIER was performed at 95 °C for 64 min using CC1 (Roche, Cat# 950-224). 2 drops of the mRNA inhibitor from mRNA Red detection kit (Roche, Cat# 760-234) were applied to the slide and incubated for 12 min. Sections were subsequently exposed for 32 min at 40°C to RNA VS PretreatPro (ACD/Bio-Techne, Cat# 322031). Before testing the target probe, RNAscope assay performance was verified for every case using an RNAscope™ 2.5 VS Positive Control Probe-Hs-PPIB (ACD/Bio-Techne, Cat# 313909) and an RNAscope™ 2.5 VS Negative Control Probe- Hs-DapB (ACD/Bio-Techne, Cat# 312039).

Hybridization was carried out with RNAscope 2.5 VS Probe-Hs-GPC5 (ACD/Bio-Techne, Cat# 521731) for 2 h at 43 °C. After hybridization, signal amplification was performed using the RNAscope VS Universal AP standard reagents (ACD/Bio-Techne, Cat# 322040) through AMP 1-AMP 7 for 32 min at 39°C per AMP, following the manufacturer’s automated sequence. Signal development was performed using the mRNA Red detection kit (Roche, Cat# 760-234) with sequential application of ACD activator for 8 min, ACD naphthol for 4 min and ACD Fast red for 12 min. After completion of RNAscope ISH, RNAscope enzyme activity was quenched using RNAscope VS CoDetectPro (ACD/Bio-Techne, Cat #322032) at 40°C for 32 min prior to protein detection. Duplex combinations included GPC5 RNAscope probe with IHC for anti-GFAP, ALDH1L1, NeuN, pS396, AT8 or 4G8, with chromogenic visualization using DISCOVERY Teal HRP Kit, as described above. Antibody concentrations are provided in Supplementary Table 2. Slides were counterstained with haematoxylin II (Roche, Cat# 790-2208) for 8 min and bluing reagent (Roche, Cat# 760-2037) for 4 min, then rinsed with Reaction Buffer and air-dried. Mounting was performed with an EcoMountTM Mounting Medium (ACD/Bio-Techne, Cat# 320409).

### Brightfield Microscopy

Brightfield imaging of chromogenic stained FFPE sections was conducted using a Leica DM2000 LED microscope equipped with 5x, 10x, 20x, 40x and 63x objectives, along with a Leica DMC2900 camera (Leica Microsystems). Additionally, high-throughput whole-slide scanning was performed using the IntelliSite Ultra Fast Scanner (Philips). In certain analysis, if needed, images were also acquired using the HALO® image analysis platform (Indica Labs, version 3.6).

### Digital Pathology

Analysis of cIHC images were performed using the HALO® image analysis platform (Indica Labs, version 4). The areas of the hippocampus, which was manually segmented into five subregions (dentate gyrus (DG), Cornu Ammonis (CA) 4, CA3, CA1/CA2, and subiculum as well as the parahippocampal cortex (PHC)) were measured. GFAP, AQP4, ALDH1L1, ALDH7A1 and GPC5-positive areas were quantified using the HALO® area quantification module. Comparisons of stained area percentages were conducted using Wilcoxon test with R Studio software.

### R2 Genomics platform database analysis

The *GFAP* and *GPC5* genes, which showed a positive correlation, were extracted from the R2 Genomics platform (https://r2.amc.nl/) from a dataset (n=173) of cognitively unimpaired individuals in the Alzheimer’s Disease Research Center (Brain-ADRC) Cotman (253 - MAS5.0 - u133p2) regrouping four brain regions: the hippocampus, the entorhinal cortex, the post-central gyrus and the superior frontal gyrus (<u>GSE48350</u>). To further refine astrocyte-relevant targets, we integrated data from the Brain RNA-seq (https://brainrnaseq.org/) and Human Protein Atlas (https://www.proteinatlas.org/) databases, selecting genes associated with astrocyte identity or function. The 500 most highly correlated genes for *gfap* or *gpc5* were then subjected to STRING pathway enrichment analysis (https://string-db.org), with the four most prevalent Kyoto Encyclopedia of Genes and Genomes (KEGG) pathways reported graphically.

## RESULTS

### Glypican 5 immunoreactivity identifies selective astrocytic and neuronal population in the human hippocampus and PHC

First, we analysed GPC5 protein distribution by cIHC in the hippocampal subregions and PHC of post-mortem control (CTL) samples (mean age at death: 89 years, 5 females, 3 males). GPC5 exhibited a discrete and spatially restricted expression pattern throughout the hippocampus (Fig. 1A). GPC5-positive astrocytes were detected in the DG and in the CA4 region but GPC5 staining appeared more diffuse in the molecular layer of the DG (Fig. 1B). GPC5 displayed a laminar distribution across the CA subfields. Strongly stained astrocytes arboring a complex and highly branched morphology were mainly found in the molecular and stratum oriens layers as well as in the subiculum (Fig. 1C–E). Strong but diffuse GPC5 immunoreactivity was observed in the neuropil of the outer two-thirds of the molecular layer (SM 2/3) of the DG in all samples. This area is enriched in vesicular GABA transporter (VGAT)-positive synapses while the inner molecular layer is enriched in vesicular glutamate transporter 1 (VGLUT1) (Fig. 1F-H). Additionally, some CA1 pyramidal neurons were variably GPC5-positive across samples (Fig. 1I). These neurons were frequently found at the two borders of the CA1, close to both the CA2 and the subiculum, whereas they were less frequent in the central CA1. The distribution of GPC5 within the subiculum and PHC was broadly similar, with an enrichment of staining in astrocytes at the glia limitans and in the upper and deeper layers as well as in some subsets of neurons (Supplementary Fig. 1A). In some CTL samples with low ABC scores, we noticed GPC5 plaque- and tangle-like staining.

**Figure 1:**
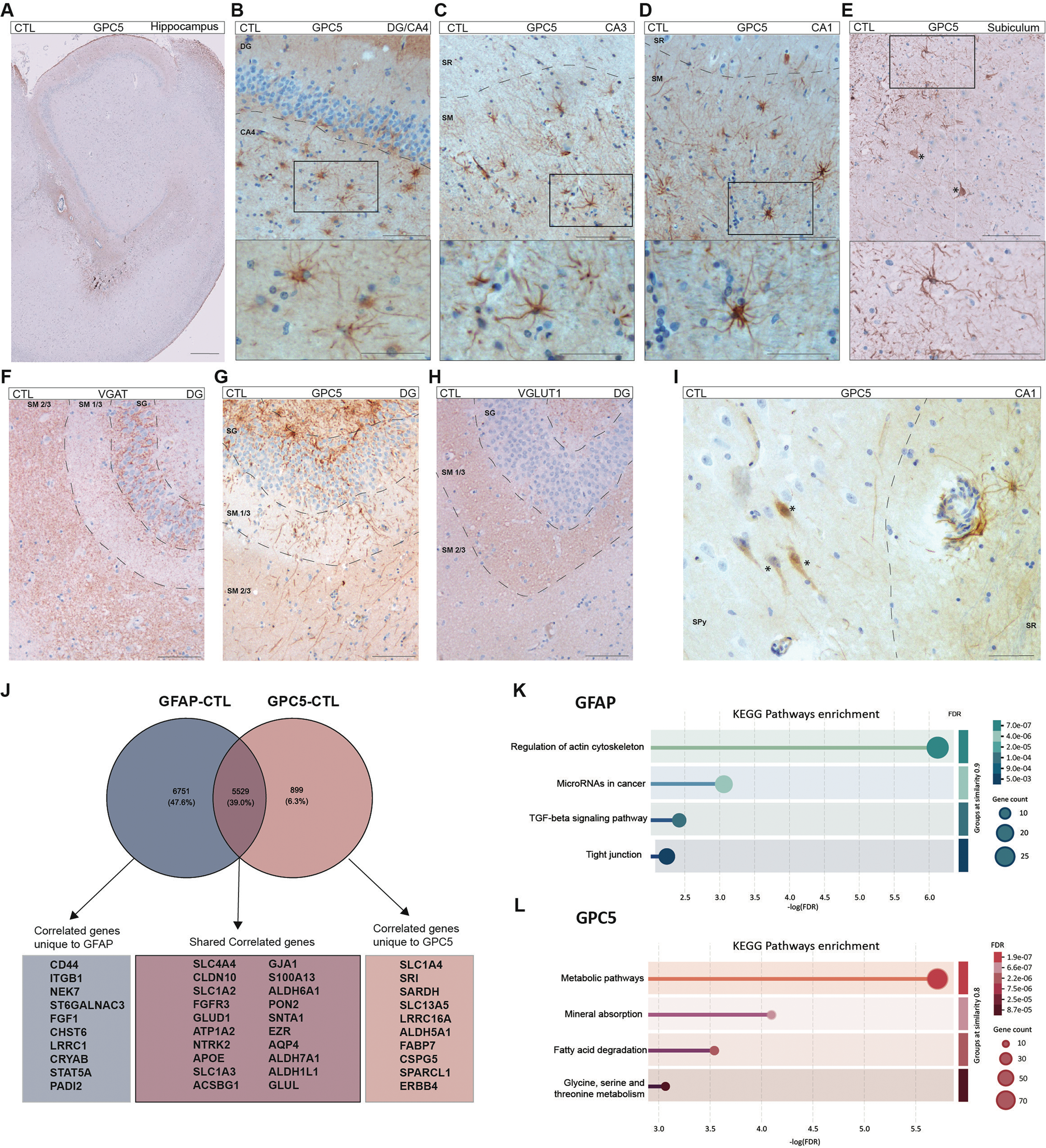
GPC5 immunoreactivity is spatially restricted in the hippocampus and associated with astrocytes and neurons. A-E. cIHC reveals the distribution of GPC5 by astrocytes in the healthy hippocampus. **A** GPC5 (DAB, brown) immunoreactivity is enriched in specific layers of the hippocampus subfields (case #5). **B** GPC5 labels many well-defined astrocytes of the CA4 but is diffuse in the DG (case #6). **C** GPC5+ astrocytes of the CA3 are mainly observed in the stratum moleculare (case #6). **D** In the CA1, the radiatum and pyramidal layers showed low density of GPC5+ astrocytes compared to the stratum moleculare (case #6). **E** In the subiculum, GPC5 labels many astrocytic branches and stains a few neurons (asterisk) (case #12). **F-H:** Representative IHC staining (DAB, brown) of (**F**) VGAT, (**G**) GPC5 and (**H**) VGLUT1 in the CTL DG show an enrichment of GPC5 in VGAT positive areas (case #2). **I** In the CA1, some few pyramidal neurons were labelled by GPC5 (asterisk) (case #6). **J-L** *GFAP* and *GPC5*-associated signatures in the R2 human brain bulk RNA-seq database. Gene expression positively correlated (p value cutoff: 0.05 FDR) with *GFAP* and *GPC5* were extracted from the <u>GSE48350</u> dataset (173 normal brain samples, 4 brain regions) in the R2: Genomics analysis and visualization platform. **J** The Venn diagram shows a shared astrocytic signature between *GFAP* and *GPC5* correlates as well as unique associations. **K-L** Functional signatures associated with (**K**) *GFAP* or (**L**) *GPC5*. The top 500 genes with the strongest positive correlation with *GFAP* or *GPC5* revealed specific KEGG pathway enrichment. **SG**: stratum granulosum; **SM 1/3**: inner stratum moleculare; **SM 2/3**: outer stratum moleculare. **SM**: stratum moleculare; **SR**: stratum radiatum. Scale bars: **A** 500 μm; **B-E** 100 μm (upper panel) 50 μm (lower panel); **F-H** 100 μm; **I** 50 μm.

### GPC5 labels specific regional astrocyte subsets in the human hippocampus and PHC

To explore the potential molecular properties of GPC5 in astrocytes, we identified genes positively correlated with either *GFAP,* a common marker for astrocytes, or *GPC5* across bulk RNA sequencing of four brain regions (hippocampus, entorhinal cortex, post-central gyrus, and superior frontal gyrus) in cognitively unimpaired individuals (CTL) from the Alzheimer’s Disease Research Center (Brain-ADRC) Cotman dataset (n=173) using the R2 Genomics platform. This analysis revealed both marker-specific gene sets and a core set of genes that were correlated with the expression of both *GFAP* and *GPC5* (Fig. 1J). *GFAP*-relevant genes included *CD44* and *ST6GALNAC3*, while *SLC1A4* and *ERBB4* were associated with *GPC5*. A set of canonical astrocytic genes; *SLC1A2*, *SLC1A3*, *ALDH1L1*, *AQP4*, *ALDH7A1*, and *APOE;* was shared between both groups, supporting the hypothesis of a common core identity. To infer biological differences, we subjected the first 500 positively correlated genes of each group to STRING pathway enrichment analysis (https://string-db.org). *GFAP*-associated genes were enriched for KEGG pathways including regulation of the actin cytoskeleton and TGF-β signalling, while *GPC5-*correlated genes showed enrichment in fatty acid degradation and glycine, serine and threonine metabolism (Fig. 1K-L).

To further characterise the identity of GPC5-positive astrocytes within the human hippocampus, we first analysed the spatial distribution patterns of three canonical and one emerging astrocytic marker: GFAP, AQP4, ALDH1L1 and ALDH7A1, in the hippocampus of our CTL cases. While GFAP, AQP4, ALDH1L1 and ALDH7A1 were all expressed throughout the hippocampus and PHC, each marker revealed distinct astrocyte features (Fig. 2 A-C; Supplementary Fig. 1B-D; Supplementary Fig. 2). In brief, GFAP staining revealed a heterogeneous distribution pattern across the subfields and layers of the hippocampus (Fig. 2Aa-d). The highest levels of GFAP staining were observed in the oriens, molecular and radiatum layers within the CA and subiculum regions. However, it was sparse and patchy in the pyramidal layer, with some areas being completely devoid of GFAP-positive astrocytes. AQP4 staining showed higher intensity in the DG and CA4, and denser labeling in the molecular layer of CA compared to the radiatum and pyramidal layers (Fig. 2Be-h). ALDH1L1 and ALDH7A1 staining exhibited a broader expression than GFAP and AQP4 throughout the hippocampus, with a uniform astrocyte density across subfields (Fig. 2Ci-l; Supplementary Fig. 2). As GPC5 distribution pattern did not match with any of these markers but partially overlapped, we performed multiplex cIHC of GPC5 in combination with GFAP (n=6), AQP4 (n=3), ALDH1L1 (n=5) or ALDH7A1 (n=4) along multiplex cIHC combining GFAP with AQP4 (n=3) or ALDH1L1 (n=5) or ALDH7A1 (n=5) in post-mortem CTL samples (Fig. 2D-H, Supplementary Fig. 3, Supplementary Fig. 4). Duplex staining with GPC5 and GFAP revealed a continuum of astrocytic phenotypes, including GPC5-only, GFAP-only, and GPC5/GFAP double-positive cells. Notably, GPC5-positive astrocytes displayed a wide range of GFAP expression levels, from GPC5/GFAP_LOW_ to GPC5/GFAP_HIGH_, predominantly in CA4 and the CA molecular layer, but also in the glia limitans and the first layer of the PHC (Fig. 2D-E, Supplementary Fig. 4C). Co-staining for GPC5 and AQP4 revealed the presence of double-positive AQP4+/GPC5+ astrocytes in the CA4 region and in the stratum oriens of the CA, and a small number in the molecular layer of the CA (Fig. 2F-G). We also observed double-positive AQP4/GPC5 cells in the second layer of the PHC with many single AQP4-positive astrocytes (Supplementary Fig. 4D). All GPC5-positive astrocytes co-expressed ALDH1L1 or ALDH7A1,thereby the two latter being considered as boarder markers than GFAP or AQP4 (Fig. 2H, Supplementary Fig. 4E). Notably, duplex staining combining GFAP, AQP4, ALDH1L1, and ALDH7A1 revealed astrocyte heterogeneity driven primarily by differential GFAP and AQP4 immunoreactivity (Supplementary Fig. 3A-C, Supplementary Fig. 4A-B). In contrast, ALDH1L1 and ALDH7A1 were broader astrocytic markers showing a staining of the entire hippocampal and PHC astrocyte population (Supplementary Fig. 3B-D; Supplementary Fig.4 B,E).

**Figure 2:**
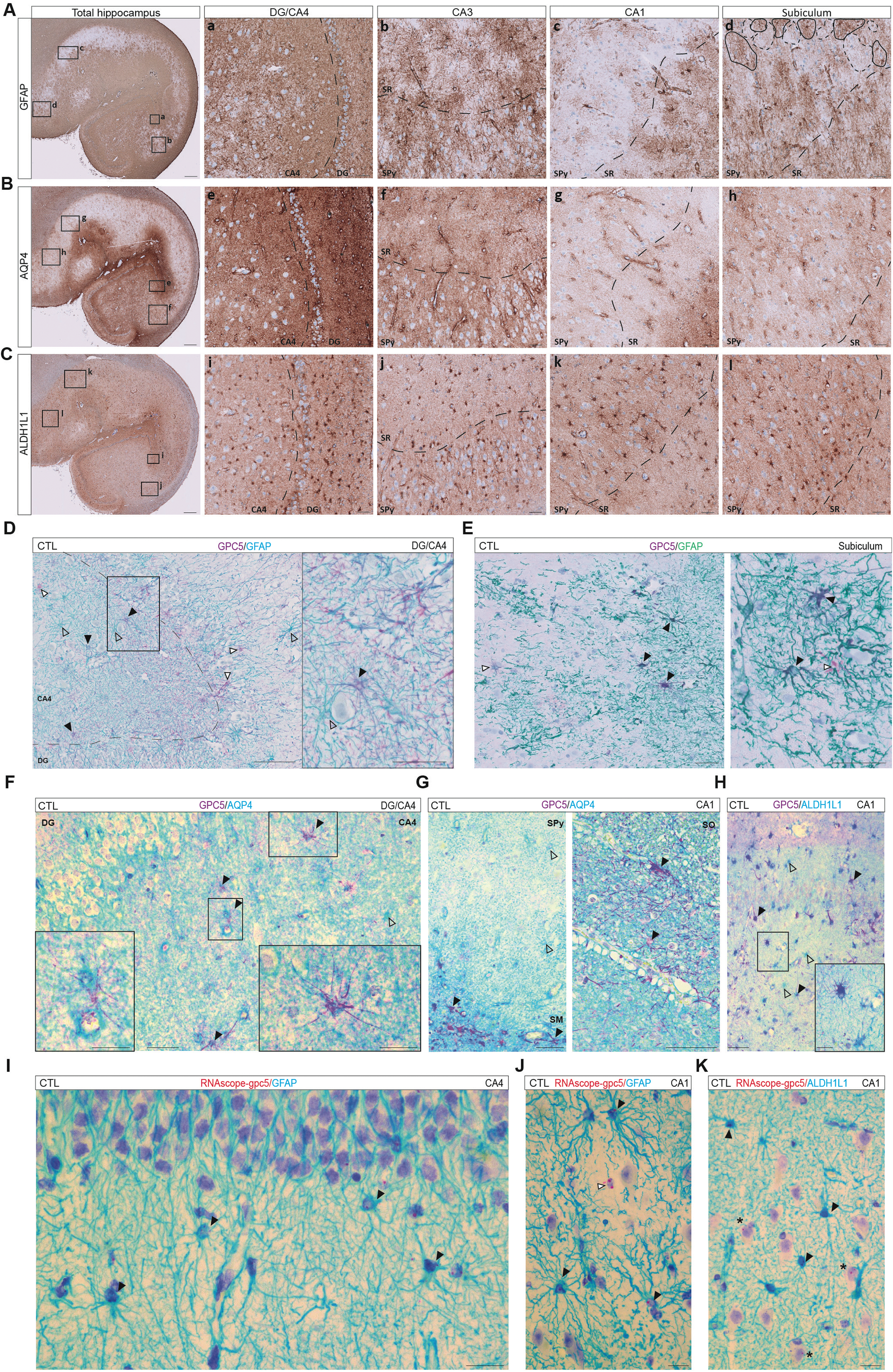
Relationship between GPC5 expression and astrocyte markers GFAP, AQP4 and ALDH1L1 across CTL hippocampi. Representative IHC of (**A**) GFAP, (**B**) AQP4 and (**C**) ALDH1L1 distribution in non-demented CTL hippocampus (case #8). **A** GFAP distribution (DAB, brown) differs between hippocampal subfields with a strong density of GFAP-positive astrocytes in (**Aa**) DG and CA4. GFAP-positive astrocytes in the (**Ab**) CA3, (**Ac**) CA1 and (**Ad**) subiculum shows a patchy distribution, especially in the pyramidal layer. In the subiculum, we distinguished two types of GFAP-positive astrocytes: cells with high levels of GFAP (GFAP_HIGH_, full lines) and cells with low GFAP expression (GFAP_LOW_, dashed lines). **B** AQP4 (DAB, brown) is highly associated with astrocytes of the (**Be**) DG/CA4, (**Bf**) CA3 and stratum moleculare of the CA compared to (**Bg**) CA1 and (**Bh**) subiculum. **C** ALDH1L1 (DAB, brown) is widely expressed by hippocampal astrocytes across all hippocampal subfields (**Ci-l**). **D-E** Co-staining of GFAP and GPC5 highlights single-positive astrocytes for either GFAP (teal, empty arrows) or GPC5 (purple, white arrows). It also reveals double-positive GFAP/GPC5 astrocytes (full arrows) (cases #2 and #9). **F-G** Co-staining of GPC5 (purple) and AQP4 (teal) revealed double-positive GPC5/AQP4 cells (dark blue, full arrows) and single AQP4 astrocytes (teal, empty arrows) in CTL DG/CA4 and CA1(case #9). **H** GPC5 staining is associated with ALDH1L1-positive astrocytes (dark blue, full arrows), a large proportion of which are negative for GPC5 (teal, empty arrows) (case #12). **I-J** cISH with RNAscope probe for *GPC5* in combination with IHC of GFAP (teal) revealed GFAP-positive astrocyte expressing *GPC5* (red dots, full arrows) in DG/CA4 and in CA1 along GFAP negative cells (white arrows) (case #15). **K** cISH with RNAscope probe for *GPC5* in combination with IHC of ALDH1L1 (teal) revealed ALDH1L1-positive astrocyte expressing GPC5 (red dots, full arrows) alongside large CA1 pyramidal neurons positive for GPC5 RNA (asterisk) (case #14). **DG**, dentate gyrus; **SM**, stratum moleculare; **SO**, stratum oriens; **SPy**, stratum pyramidale; **SR**, stratum radiatum. Scale bars: **A-C**: 500 μm (low magnification), 50 μm (high magnification **a–l**);**D**: 100 μm (low magnification), 50 μm (high magnification**); E**: 50 μm and 20 μm (right panel); **F**: 50 μm; **G**: 50 μm and 20 μm (right panels); **H**: 50 μm and 20 μm (high magnification); **I-K**: 20 μm.

To further validate the expression of *GPC5* in an anatomically selective population of astrocytes, we combined chromogenic RNAscope ISH with cell-type identification by cIHC. We used the RNAscope *GPC5* human probe (red dots) alongside GFAP or ALDH1L1 staining (teal) (Fig. 2I-K). *GPC5* probe revealed the presence of *GPC5*-expressing, GFAP-positive and ALDH1L1-positive astrocytes, consistent with their anatomical distribution as observed by IHC. Some CA1 neurons were also found to be positive for the *GPC5* probe (Fig. 2K). These observations highlighted the molecular heterogeneity of hippocampal astrocytes and GPC5 as a marker of a specific sub-regional astrocyte subpopulation.

### GPC5 is restricted to specific pyramidal neuron populations in the human hippocampus and PHC

We then investigated the distribution of neuronal GPC5 in more detail using cIHC, combining the neuronal marker NeuN with GPC5 in CTL samples (n=3). In the hippocampus and PHC, most NeuN-positive neurons were GPC5-negative (Fig. 3A). However, a small proportion in the pyramidal layer of CA1 were double-positive, in line with our observations from single staining (Fig. 3B). To determine whether GPC5 neuronal staining could be due to an astrocyte-secreted source of GPC5, we employed RNAscope *GPC5* human probes in conjunction with NeuN cIHC. The RNAscope *GPC5* probe labelled NeuN-positive neurons in the periphery of the CA1 (in close proximity to the CA2 region and the subiculum) and the PHC, whereas the granular neurons of the DG were negative (Fig. 3C-E). This indicates that GPC5 is also expressed by a subset of hippocampal neurons.

**Figure 3:**
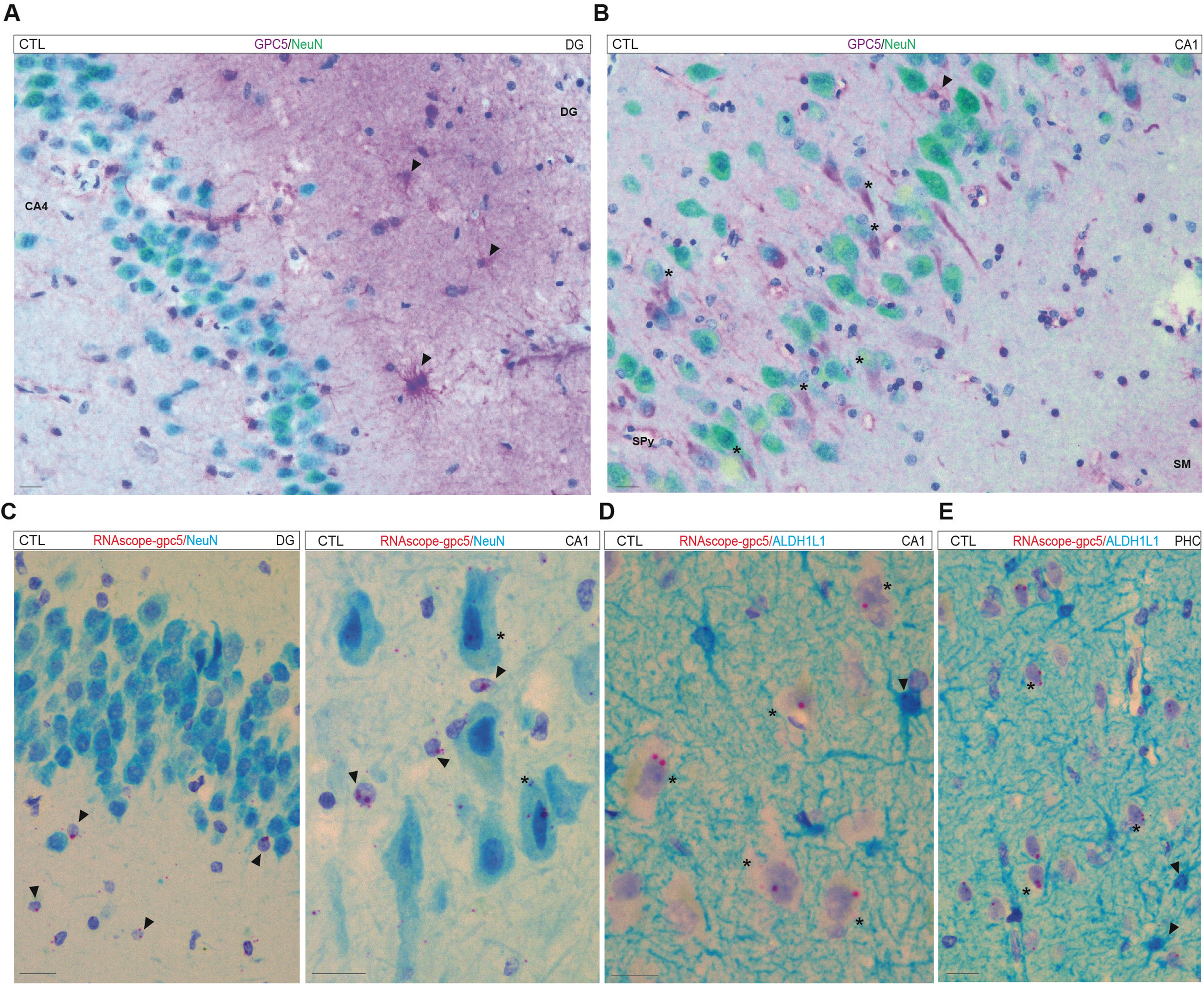
A subset of pyramidal neurons expresses GPC5 at the RNA and protein levels in the CTL hippocampus. A-B. Co-staining with NeuN (green) and GPC5 (purple) shows that GPC5 is not present in the granular neurons of the (**A**) and is only found in certain subgroups of CA1 pyramidal neurons (asterisk, **B**). The proportion of GPC5 astrocytes is higher in the CA4, DG and SM of CA1. They are sparse in the Spy of the CA1 (full arrows) (case #8). **C** cISH of *GPC5* RNAscope probe, in combination with IHC of NeuN (teal), confirmed the absence of *GPC5* mRNA in granular neurons, and its presence in a small proportion of CA1 neurons (asterisk). By contrast, most of the cells expressing *GPC5* were similar in size to astrocytes (full arrows) (case #13). **D** cISH of *GPC5* RNAscope probe in combination with IHC of ALDH1L1 (teal) revealed ALDH1L1-positive astrocyte expressing *GPC5* (red dots, full arrows) alongside large CA1 (**D**) or PHC (**E**) pyramidal neurons positive for *GPC5* RNA (asterisk) (case #14). Scale bars: **A-E**: 20 μm.

### GPC5 and canonical astrocyte markers are differently affected in AD and PDD

To investigate disease-associated alterations in astrocyte subtypes, we performed cIHC and quantified DAB-stained area for GFAP, AQP4, ALDH1L1, ALDH7A1 and GPC5 in post-mortem hippocampal sections from individuals with AD (n=9, average age at death: 73.4 years, 6 females, 3 males) and PDD (GFAP, AQP4, ALDH1L1 and GPC5: n=10, average age at death: 74.6 years, 5 females, 5 males; ALDH7A1: n=8, average age at death: 73.8 years, 3 females, 5 males), compared to CTLs. All the tested markers followed a specific pattern across disease groups and hippocampal subfields (Fig. 4A-O). GFAP-stained area was significantly reduced in AD relative to CTLs and PDD in DG, CA4 and CA3. In CA1/CA2, GFAP intensity was decreased in both AD (p value=0.07) and PDD (p value=0.04) cases. Despite the reduced staining, we observed focal areas of intense GFAP expression. In some of these areas in AD, GFAP-positive astrocytes displayed a hypertrophic appearance polarised towards presumed amyloid-β (Aβ) plaques, especially in CA1. We also found clusters of astrocytes expressing high level of GFAP in PDD (Fig. 4A-C). AQP4-postive area, however, was significantly increased in AD compared to both CTLs and PDD in CA3, CA1/CA2 and subiculum. In PDD, the distribution of AQP4 was comparable to CTL cases to the exception of CA4 (p value=0.03) showing significant lower levels (Fig. 4D-F). Stained area of ALDH1L1 was significantly decreased in both AD and PDD. In AD cases, the decrease was more prominent in DG, CA4, CA3 and PHC, whereas in PDD the decrease was significant for all subfields and PHC. In both AD and PDD, we observed a general reduction of ALDH1L1 intensity in the astrocyte arborisation (Fig. 4G-I). ALDH7A1 followed the general ALDH1L1 pattern, showing a significant and strong reduction in PDD cases globally, with the strongest decrease seen in the CA3, CA1/CA2, subiculum and PHC subregions. There was a significant decrease in the subiculum and PHC of AD cases (Fig. 4J-L). GPC5 displayed a specific pattern, remaining globally stable in stained area across diseases. Besides, we observed a significant decrease of GPC5 staining area in DG of AD patients (p value=0.02 for CTL vs AD, p value=0.03 for AD vs PDD), which was associated with the loss of staining in the molecular layer. Nevertheless, GPC5-positive astrocytes exhibited a partially altered spatial organisation in AD. They appeared more scattered and forming small clusters across the hippocampus. GPC5-positive astrocytes adopted a range of morphologies, ranging from thin to enlarged. Furthermore, we observed plaque-like morphologies labelled by a more diffuse GPC5 staining, which was often associated with polarised GPC5-positive astrocytes surrounding the plaques (Fig. 4M-O; Fig. 5A-C). We also found a stronger neuronal GPC5 staining in the form of tangle-like and ghost-tangle like structure (Fig. 5C).

**Figure 4:**
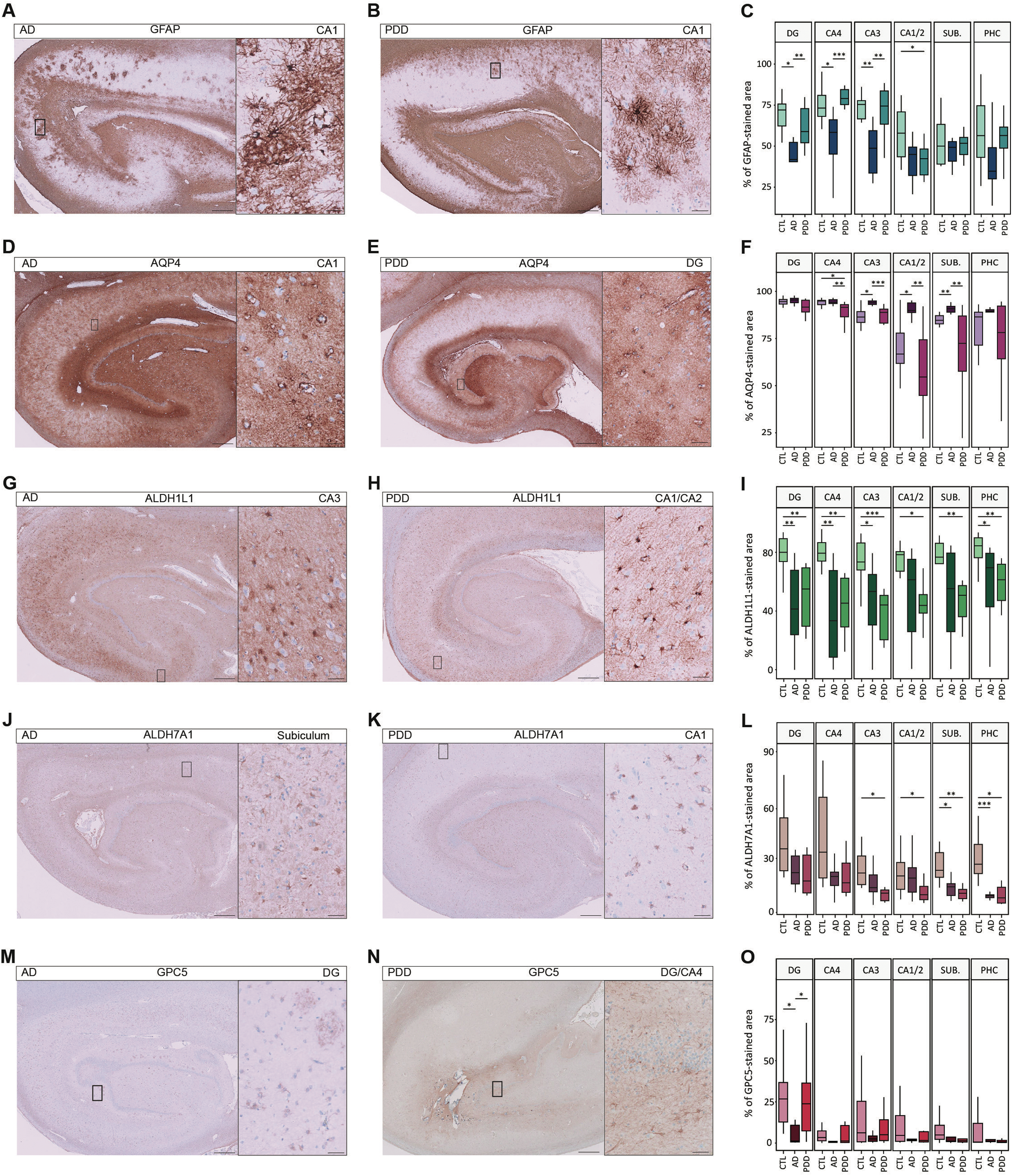
Differential alterations of astrocytic markers in AD and PDD. A-B. GFAP-positive astrocytes (DAB, brown) display a dysmorphic morphology in AD and PDD hippocampi and form cluster of cells reminiscent of reactive glial nets in AD (**A**) and reactive gliosis in PDD (**B**) (AD case #34; PDD case #51). **C** The quantitative analysis of GFAP positive area in the different hippocampal subfields of patients with AD and PDD (surface coverage area of the DAB staining) shows a strong decrease of GFAP in the DG, CA4 and CA3 of AD cases, however only in the CA1/CA2 of PDD patients (n=8 CTL, n=9 AD, n=11 PDD). **D-F** AQP4 immunoreactivity (DAB, brown) is high in astrocytes in both AD and PDD and is significantly increased in the CA3, CA1/CA2 and subiculum of patients with AD (AD case #30; PDD case #42) (n=8 CTL, n=9 AD, n=10 PDD). **G-I** ALDH1L1 staining (DAB, brown) is generally decreased among astrocytes as compared to CTLs, and present less complex arborisations. Its stained area is mainly affected in the DG, CA4 and CA3 of AD cases and overall decreased in all hippocampal subfields in PDD (AD case #30; PDD case #43) (n=8 CTL, n=9 AD, n=10 PDD). **J-L** ALDH7A1 immunoreactivity (DAB, brown) is decreased in disease notably in the CA3, CA1/CA2 and PHC in PDD and in the subiculum in both AD and PDD patients (AD case #29; PDD case #42) (n=8 CTL, n=9 AD, n=8 PDD). **M-O** GPC5 density (DAB, brown) remains stable across disease conditions. **M** In AD, GPC5 labeling reveals GPC5 concentrated in potential Aβ plaques (zoom-in). **N** In PDD, the distribution pattern of GPC5+ is similar to that of the CTL but it also shows enlarged and potentially reactive astrocytes (zoom-in) (AD case #29; PDD case #48) (n=8 CTL, n=9 AD, n=10 PDD). Statistical analysis: Wilcoxon test, * *p*<0.05, *\*\* p<0.01, *** p<0.001*. Scale bars: low magnification: 500 μm; high magnification: 50 μm.

**Figure 5:**
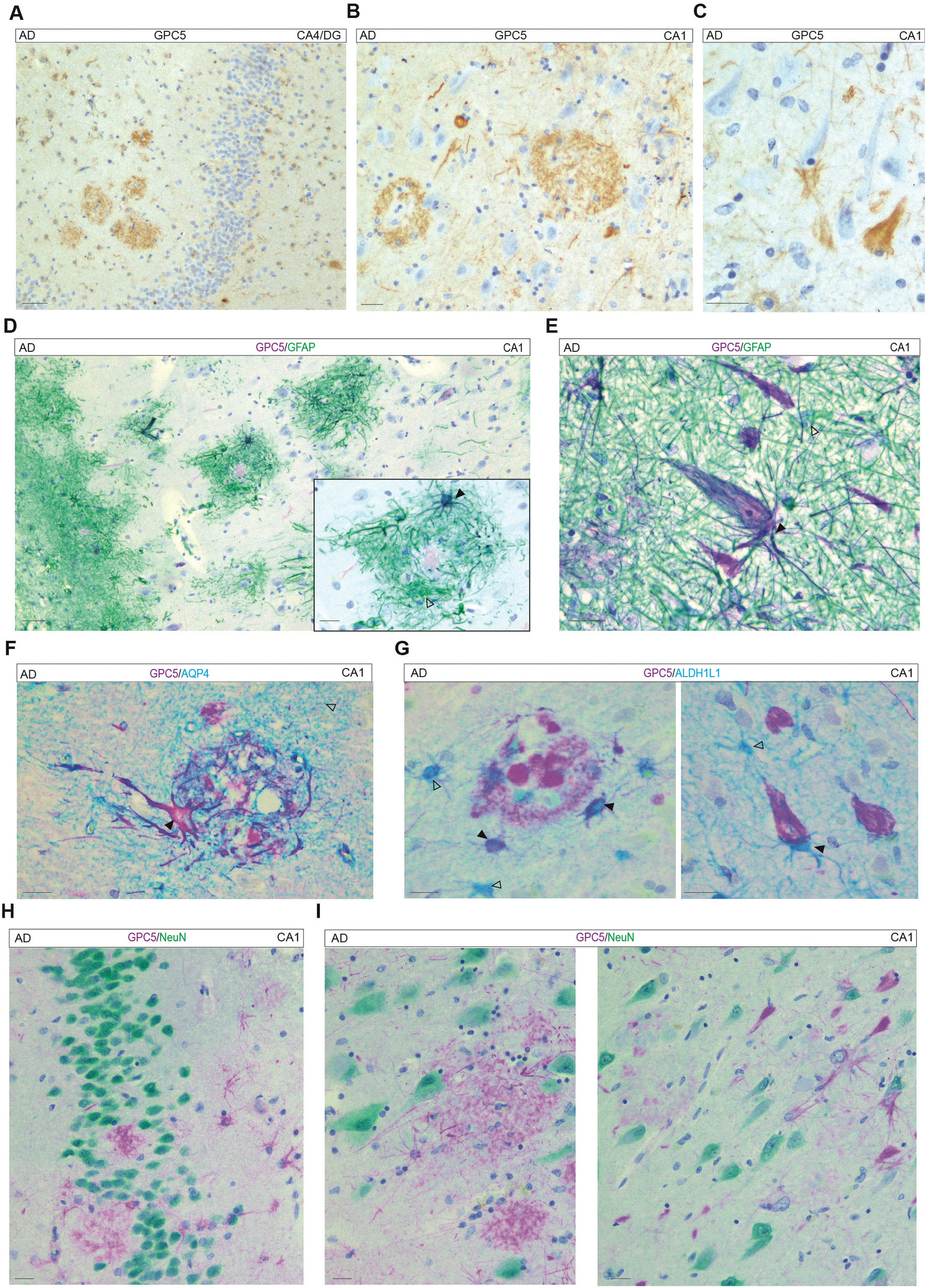
GPC5 staining is affected in patients with AD. A-C. GPC5 staining is altered in AD. GPC5 staining (DAB, brown) is associated with astrocytes, neurons but also Aβ plaques (**A-B**) and tangles (**C**) (case #28 and 29). **D-G** Multiplex cIHC of GPC5 (purple) with (**D-E**) GFAP (green); (**F**) AQP4 (teal) or (**G**) ALDH1L1 (teal) reveal a heterogeneity of GPC5 signatures within astrocytes. GPC5 is associated with some astrocytes that surround Aβ plaques stained with GFAP, AQP4 or ALDH1L1 as well as in the plaque cores. GPC5 is also found in some astrocytes in close proximity to GPC5-positive tangles, though not systematically (case #34 (D,F,G); case #28 (E); case #33 (F)). **H-I** Co-staining of GPC5 (purple) with NeuN (green) further revealed that GPC5 immunoreactivity was associated with astrocytes and with Aβ plaque-like and neuronal tangle-like structures. White and black arrows indicate single and double-positive astrocytes, respectively. Scale bars: **A**: 50 μm; **B-I**: 20 μm.

To further characterise the effect of neurodegeneration on GPC5-astrocyte identity, we next used multiplexed cIHC to co-stain GPC5 with astrocyte markers (n=6 GPC5/GFAP; n=3 GPC5/AQP4; n=6 GPC5/ALDH1L1; n=6 GPC5/ALDH7A1) in AD hippocampus and PHC (Fig. 5D-G, Supplementary Fig. 5). By combining GPC5 with GFAP, we observed a variety of profiles, a high proportion of single-positive GFAP, some double-positive GFAP/GPC5 cells displaying intensity of GFAP ranging from low to high; and a small number of single-positive GPC5 cells. GPC5/GFAP cells exhibited various morphologies, ranging from very elongated to enlarged. In both the hippocampus and the PHC, GPC5+ astrocytes that formed nets around GPC5-positive Aβ plaque-like structures were all positive for GFAP but were less frequent than single positive GFAP astrocytes (Fig. 5D, Supplementary Fig. 5A). GPC5 staining also revealed tangle-like structures near GPC5-positive astrocytes, often double stained by GFAP forming an astrocyte tangle mesh, especially in the CA1 and CA2 (Fig. 5E). Multiplexed cIHC of GPC5 with AQP4, ALDH1L1 or ALDH7A1 in the AD hippocampus and PHC revealed plaques surrounded by a subset of double-positive astrocytes and showed, frequently but not systematically, double-positive astrocytes around GPC5 positive tangle structures (Fig. 5F-G; Supplementary Fig. 5B-D). Although ALDH1L1 expression was reduced in AD, it was still present in double-positive GPC5+/ALDH1L1+ cells. The mosaic pattern of GFAP, AQP4, ALDH1L1 and ALDH7A1 immunoreactivity in the hippocampus and PHC of CTLs was largely preserved in AD (Supplementary Fig. 5E-J), although there was greater variation in morphology and intensity among GFAP/AQP4 double-positive cells (n=3 GFAP/AQP4; n=6 GFAP/ALDH1L1; n=6 GFAP/ALDH7A1). Co-staining with NeuN revealed a few GPC5/NeuN neurons, as well as GPC5-positive tangle-like structures that were not stained by NeuN (Fig. 5H-I). It also highlighted GPC5 astrocytes in close proximity to Aβ plaques and GPC5-positive tangle-like structures and the poor association of GPC5-positive neurons with Aβ plaques.

### GPC5, which is associated with A**β** plaques, is mainly expressed by astrocytes surrounding the plaques

We then examined the association between GPC5 and Aβ plaques in post-mortem AD samples by co-staining a marker of amyloid (4G8) with GPC5. Duplex staining with 4G8 (n=3) confirmed the frequent, though not systematic, association of GPC5+ astrocytes with Aβ plaques across hippocampal subfields and PHC (Fig. 6A-B). GPC5 was also found to be diffuse in the core. Triple staining with 4G8, GPC5 and GFAP (n=3) confirmed that GPC5/GFAP double-positive astrocytes are often associated with Aβ plaques, albeit in lower proportions than GFAP single-positive astrocytes, across hippocampal subfields and PHC (Fig. 6C-D). Using cISH for the *GPC5* RNAscope probe in combination with 4G8 cIHC, we found that cells compatible with astrocytes based on their nuclear size were producing *GPC5*. The larger surrounding cells, which were probably neurons, were mostly negative for *GPC5* (Fig. 6E-F).

**Figure 6:**
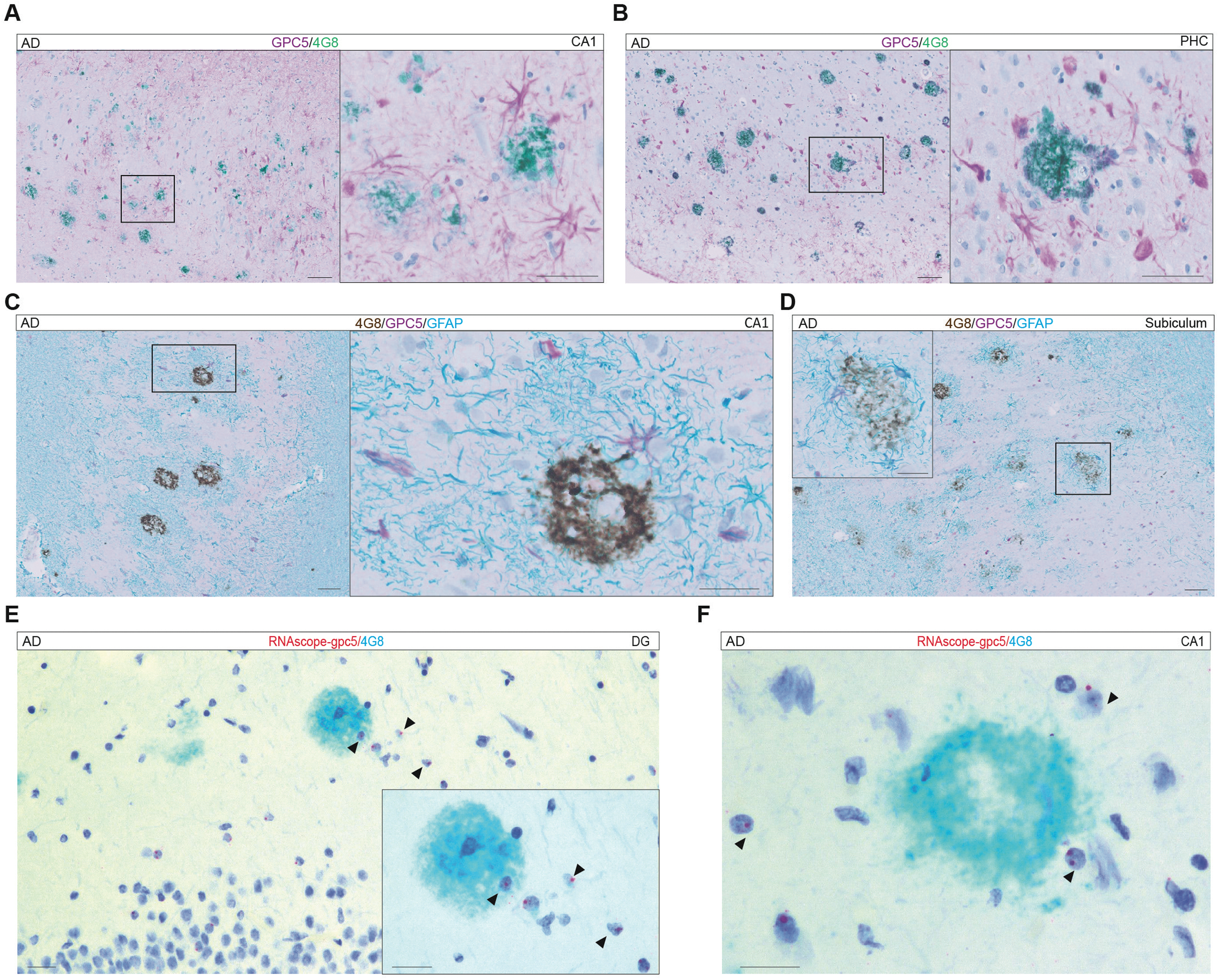
Some astrocytes associated with Aβ plaques express GPC5. A-B. Some astrocytes forming a net around Aβ plaques (4G8, green) are stained for GPC5 (purple) in the hippocampus (**A**) as well as the PHC (**B**) (cases #25, #32). **C-D** The expression of GPC5 (purple) in GFAP astrocytes (teal) surrounding Aβ plaques (DAB, brown) is heterogeneous in CA1 (**C**) or subiculum (**D**) (case #33). **E-F** cISH using an RNAscope probe for *GPC5*, in combination with IHC of 4G8 (teal) to reveal the Aβ plaques, confirmed the presence of *GPC5* mRNA in cells of a similar size to astrocytes (full arrows) in the DG and CA1 of the hippocampus (case #38). Scale bars: **A-D** low magnification: 100 μm, high magnification: 50 μm; **E-F**: 20 μm.

### GPC5 is associated with tau tangles but not with synuclein pathology

To investigate the relationship between GPC5 and tau tangles, we double-stained tissue sections for GPC5 and either AT8 (n=6) or pS396 (n=6). These are two markers of different stages of tau tangle maturation: AT8 covers a broader range, from early to advanced stages, whereas pS396 is more specific to advanced stages^32,33^. GPC5 astrocytes were frequently found near AT8-tangles. However, there was no overlap between AT8 and GPC5 staining (Fig. 7A). Double staining with pS396 revealed that pS396 tangles were frequently stained with GPC5 and were often surrounded by GPC5 astrocytes (Fig. 7B).

**Figure 7:**
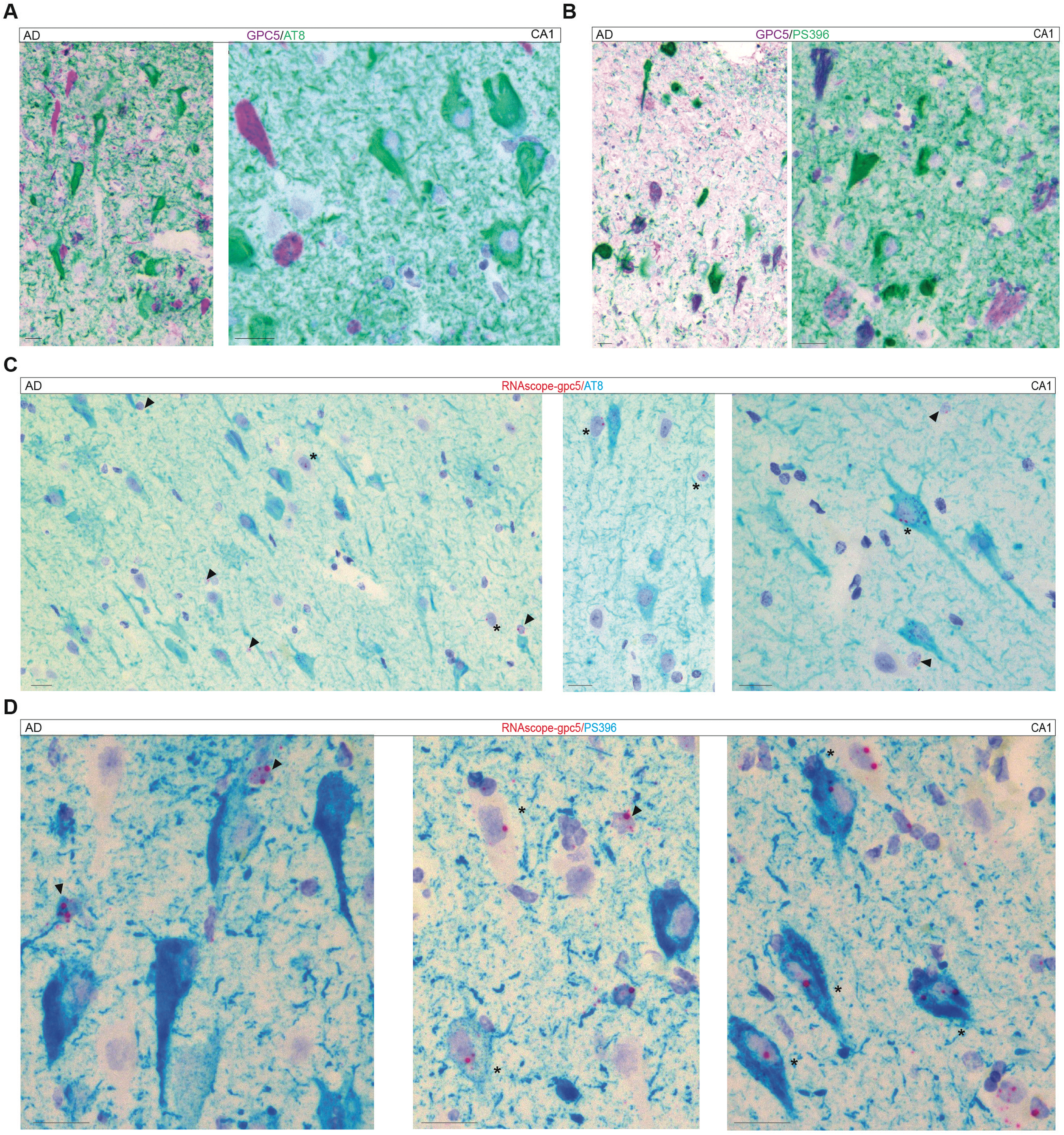
GPC5 staining, which is strongly associated with PS396-positive neurofibrillary tangles, is derived from both neuronal and astrocytic sources. **A** Tau tangles labeled by AT8 (green) show little co-localization with GPC5 (purple) (cases #37, #40). **B** PS396 (green) tangles are frequently immunoreactive for GPC5 (purple) (double positive in dark purple) (cases #37, #40).) **C** Only a small subset of neurons bearing AT8-positive neurofibrillary tangles (teal) expresses *GPC5*, as detected by RNAscope for *GPC5* (red dots) (asterisks for *GPC5*-positive neurons; full arrows for cells of a similar size to astrocytes *GPC5*-positive, case #39). **D** Combined cISH using an RNAscope probe for *GPC5* with IHC for PS396 (teal) shows that neurons bearing PS396-positive tangles frequently express *GPC5* mRNA (red dots, asterisks for *GPC5*-positive neurons; full arrows for cells of a similar size to astrocytes *GPC5*-positive, case #41). Scale bars: **A-F**: 20 μm.

Using cISH for the *GPC5* RNAscope probe in combination with cIHC of either AT8 or PS396 revealed a weak association between AT8 staining and *GPC5* RNA-producing neurons, but a stronger association with PS396 (Fig 7C-D). It should be noted that *GPC5* tangle-bearing neurons were often associated with astrocyte-like cells expressing *GPC5* (Fig 7C-D). Finally, in PDD samples stained for GPC5, GFAP and pSYN (pSYN 81A), we found no clear association between GPC5 astrocytes, Lewy bodies or neurites (Supplementary Fig. 6A). However, we observe that some dysmorphic GFAP+ astrocytes exhibited intracellular accumulation of phosphorylated α-synuclein (Supplementary Fig.6B).

## DISCUSSION

Astrocytes and neurons exhibit significant regional and molecular heterogeneity^34,35^. Nevertheless, the manner in which this diversity manifests in the human brain and its contribution to the onset or progression of NDDs remains poorly defined. In this study, we integrated cIHC and cISH^36^ with quantitative digital pathology in non-demented CTLs and NDDs such as AD and PDD. We showed that GPC5 is a marker of this heterogeneity in physiological and disease conditions. GPC5 is physiologically expressed by anatomically distinct subpopulations of astrocytes throughout the hippocampus and by a restricted population of pyramidal neurons at the CA1 border, with additional enrichment in the DG neuropil. In AD, this anatomical distribution is profoundly altered, with GPC5 redistributing toward Aβ plaques and tau pathology. In contrast, we found no evidence of an association between GPC5 and α-synuclein pathology in PDD. Together, these findings identify a disease-specific redistribution of GPC5 in AD, highlighting its potential role in hippocampal organisation and in the tissue response to AD pathology.

### GPC5 is a marker of astrocyte heterogeneity and their response to AD pathology

Although it is widely accepted that astrocytes exhibit notable differences in morphology, properties, and functionality between and within brain regions^37^, this heterogeneity is still poorly understood in the context of onset and progression of brain diseases. Astrocytes are traditionally defined by a star-shaped morphology in grey matter, but specialised subtypes exist throughout the CNS. These include the fibrous astrocytes of the white matter, the Müller glia in the retina, the Bergmann glia in the cerebellar cortex, or the interlaminar astrocytes and the varicose projection astrocytes found in the superficial cortical layers of the human brain^38^. The presence of a reactive astrocytic phenotype is commonly assessed using markers such as glial fibrillary acidic protein (GFAP), which remains the most widely used astrocyte marker^39,40^, alongside ALDH1L1, AQP4, S100 calcium-binding protein beta (S100b), or excitatory amino acid transporter 1/2 (EAAT1/2). Furthermore, GFAP in cerebrospinal fluid (CSF) and blood has also emerged as a promising biomarker for AD^41,42^. However, the extent to which it correlates with parenchymal astrocytic changes is still being debated^43^. In addition to GPC5 staining, which identifies a subset of astrocytes associated with anatomy and pathology, we also provide relevant information and a broader overview of astrocyte diversity in health and disease. GFAP does not stain all hippocampal astrocytes uniformly but labels a large subset. This subset shows a heterogeneous and subfield-specific distribution in age-matched CTL samples. Regions with low GFAP staining remained populated by ALDH1L1-, or ALDH7A1-positive, and, to a lesser extent, AQP4-positive astrocytes. This supports the notion that astrocyte diversity cannot be captured by a single marker. Consistent with observations in rodents^44,45,46,47^, our findings suggest that ALDH1L1 and ALDH7A1 (Antiquitin) are broader astrocyte markers than GFAP, capturing the majority, if not all, astrocytes.

In AD, we observed a marked reduction of GFAP stained area, particularly in hippocampal subfields that normally display the highest GFAP levels, whereas PDD showed no comparable change. Although this contrasts with mouse model reports of increased GFAP in AD^48^, the discrepancy may reflect disease stage, with astrocyte reactivity peaking early and declining during advanced neurodegeneration^49^. Notably, GFAP-positive astrocytes remained associated with Aβ plaques, supporting their involvement in plaque-associated glial responses^33^. In parallel, AQP4 was strongly upregulated in AD, consistent with impaired glymphatic function and astrocyte dysfunction^50,51,52^. By contrast, PDD was characterised by reduced expression of ALDH1L1 and ALDH7A1, suggesting astrocytic alterations specific to the disease that may affect folate metabolism, neurogenesis, and extracellular matrix homeostasis^53–59^. Overall, the distribution patterns of GFAP, AQP4, ALDH1L1, and ALDH7A1 indicate a partial-to-severe loss of astrocytic homeostatic signatures in advanced AD and, to a lesser extent, PDD.

Our analysis of GPC5 expression reveals an additional dimension of astrocyte heterogeneity that may be relevant to both normal brain function and NDDs. GPC5 belongs to the glypican family of GPI-anchored heparan sulfate proteoglycans (GPC1–GPC6) and is broadly expressed throughout the mammalian brain during development and adulthood^60^. Among these, GPC5 is notably expressed by human astrocytes and its function has been associated with thalamocortical synapse maturation and maintenance^22^. The highly selective, laminar, and subregional distribution of GPC5-positive astrocytes suggests that distinct astrocyte populations are associated with specific neuronal circuits in the adult human brain, particularly within the hippocampus and adjacent cortical regions. This pattern is markedly changed in AD but not in PDD. Notably, GPC5 expression has been reported to be dysregulated in various NDDs, including fronto-temporal dementia^24^, multiple sclerosis^25^ and AD^26,27^. This suggests a broad role of GPC5 in brain pathology. Given GPC5’s role in synapse regulation, changes in its expression could influence synaptic dysfunction and neuronal hyperexcitability^22,26^. In AD, the spatial organization of GPC5-positive astrocytes differed markedly from that observed in age-matched CTLs, particularly within CA1/CA2 and the PHC, where GPC5-positive astrocytes frequently clustered around Aβ plaques and were observed in close proximity to neurofibrillary tangles. Because GPC5 expression has been associated with homeostatic astrocyte states^18,19,61,62^, these observations raise the possibility that GPC5-expressing astrocytes represent a protective or compensatory astrocyte subtype recruited to sites of pathology. Our findings further highlight the cellular heterogeneity of reactive glial nets (RGNs)^63^, suggesting that astrocyte populations participating in these structures may vary across anatomical regions and disease stages and could support specific functions.

### AD disrupts the physiological distribution of neuronal GPC5

Our cISH analyses suggest that the source of plaque-associated GPC5 is predominantly astrocytic. However, GPC5 was also expressed by a specific group of pyramidal neurons in age-matched CTLs and was found in some tangle-bearing neurons. This suggests that there are distinct astrocytic and neuronal contributions to its distribution in AD pathology. Reduced GPC5 expression in the outer molecular layer of the dentate gyrus in AD, a region with a high concentration of VGAT-positive inhibitory synapses, may contribute to changes in the balance of excitation and inhibition in the hippocampus and, consequently, to alterations in network excitability. Additionally, we observed diffuse GPC5 immunoreactivity within Aβ plaque cores and neurofibrillary tangles, indicating that GPC5 may also become incorporated into hallmark proteinaceous lesions of AD. Glypicans such as GPC1 and GPC4 are intimately involved in the Aβ plaque formation^64,65,66^ and are found in diffuse and core plaques^67^. GPC5 have recently been found to be associated with AD brain proteome and plaques in CRDN8 AD mouse model and humans^68^. Our cISH analyses further suggest that astrocytes represent a local source of GPC5 in the vicinity of Aβ plaques. Diffuse GPC5 immunoreactivity was also observed around neurofibrillary tangles. This pattern was reminiscent of the fine GFAP-positive astrocytic processes reported to accumulate around ghost tangles; however, the expression of GPC5 in a subset of tangle-bearing neurons suggests that both astrocytic and neuronal sources may contribute to its tangle-associated localization. Previous studies have implicated another astrocyte-derived glypican, GPC4, in AD pathogenesis. GPC4, which is secreted by a subset of astrocytes, has been reported to interact with ApoE4 and to promote tau hyperphosphorylation *in vitro* and in mouse models^69^. In this context, the preferential association of GPC5 with pS396-positive rather than AT8-positive tau pathology suggests that GPC5 may be associated with specific stages of tau aggregation or with the maturation of neurofibrillary tangles.

### Limitations of the study

This study has several limitations. Firstly, our cohort included a limited number of brain samples, which may affect generalisability. Secondly, the study focused on advanced disease stages. Further validation in larger, independent cohorts is required, as well as an assessment of astrocyte and neuronal diversity in other key regions.

## Conclusion

Together, these findings identify GPC5 as a molecular signature of spatially defined astrocyte and neuronal populations and demonstrate its selective reorganisation in AD but not in PDD, supporting a role for GPC5-linked cellular domains in disease-specific vulnerability. This spatial reorganisation is accompanied by the redistribution of GPC5 towards Aβ plaques and neurofibrillary tau tangles, indicating an association with the core histopathological hallmarks of AD.

## DECLARATIONS

### Ethics approval

Use of human brain *post-mortem* samples for research was approved by the respective Ethic Panels of the Douglas-Bell Canada Brain Bank (Douglas Mental Health University Institute, Montreal, QC, Canada), the Netherlands Brain Bank (Netherlands Institute for Neuroscience, Amsterdam) and the GIE-Neuro-CEB biobank (Groupe Hospitalier Pitié-Salpêtrière, Paris, France) as well as of the University of Luxembourg (ERP 16-037 and 21-009).

### Data availability

Raw data is available on request.

### Consent for publication

All authors have consented for the publication of manuscript.

## Acknowledgments

We would like to express our deepest gratitude to the brain donors and their families for their generous support of this project. We would also like to thank the staff at the brain banks for their dedication and assistance.

## Author contributions

DSB conceived and supervised the study. FJ, MMM, and SSc performed the experiments. FJ, MMM, SSc, and GH generated and analyzed the statistical data. FJ, SSc, and DSB prepared the figures. DM, NM, and NBB provided the human brain cohort and neuropathological reports. FJ, MMM, MM, and DSB wrote the manuscript with input from all authors.

