## Supplementary figures and images for "Glypican-5 delineates spatially distinct astrocyte and neuronal populations in the human hippocampus and is selectively remodelled in Alzheimer’s disease"

### Supplementary Figure 1

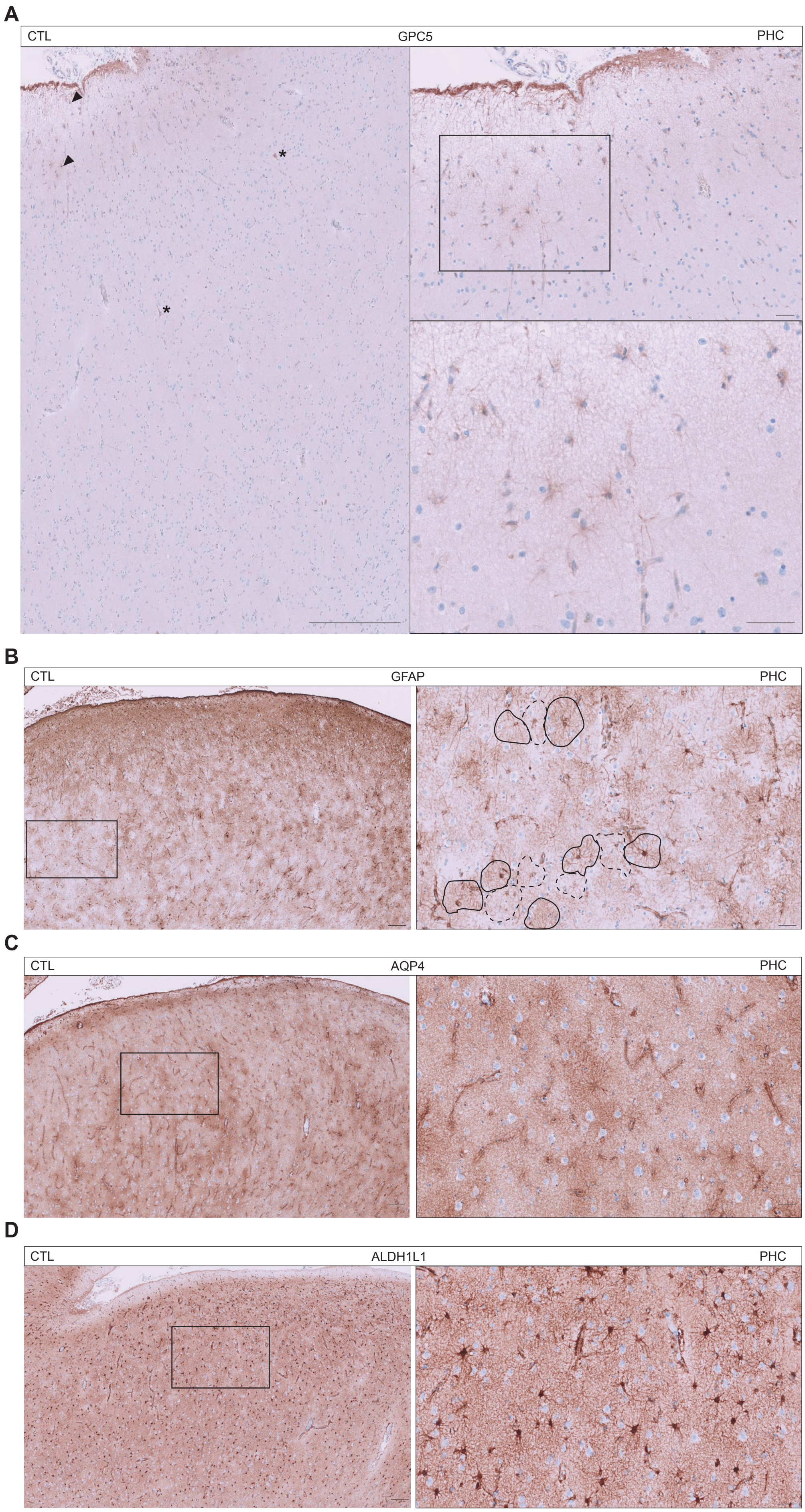


**SUPPLEMENTARY FIGURE 1**

### Supplementary Figure 2

A

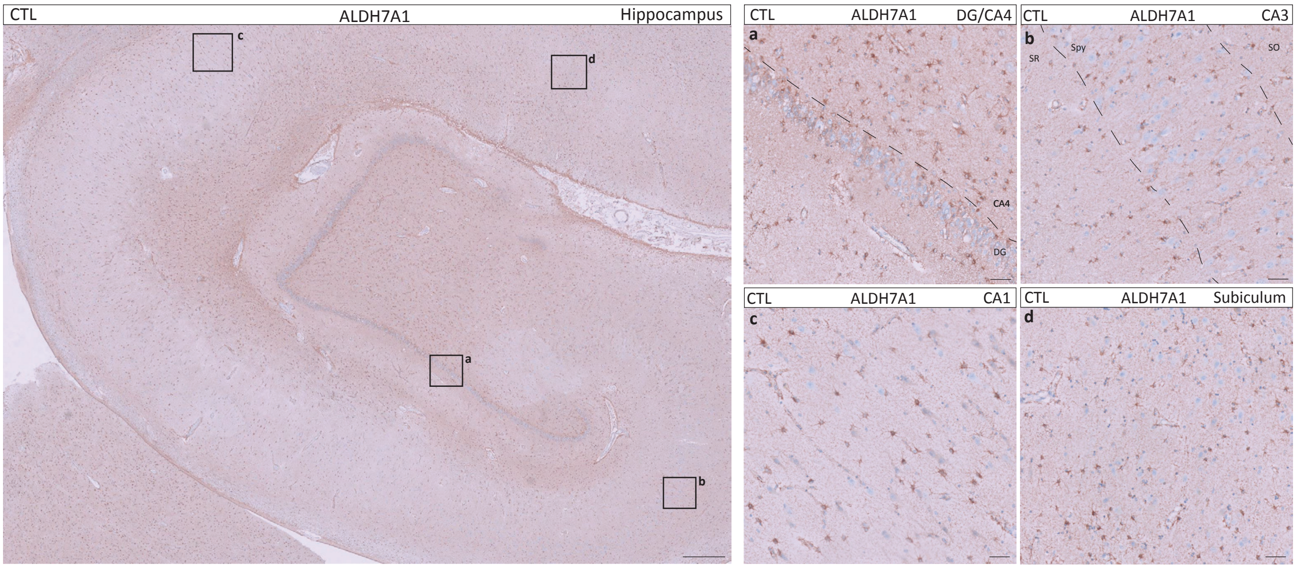


SUPPLEMENTARY FIGURE 2

### Supplementary Figure 3

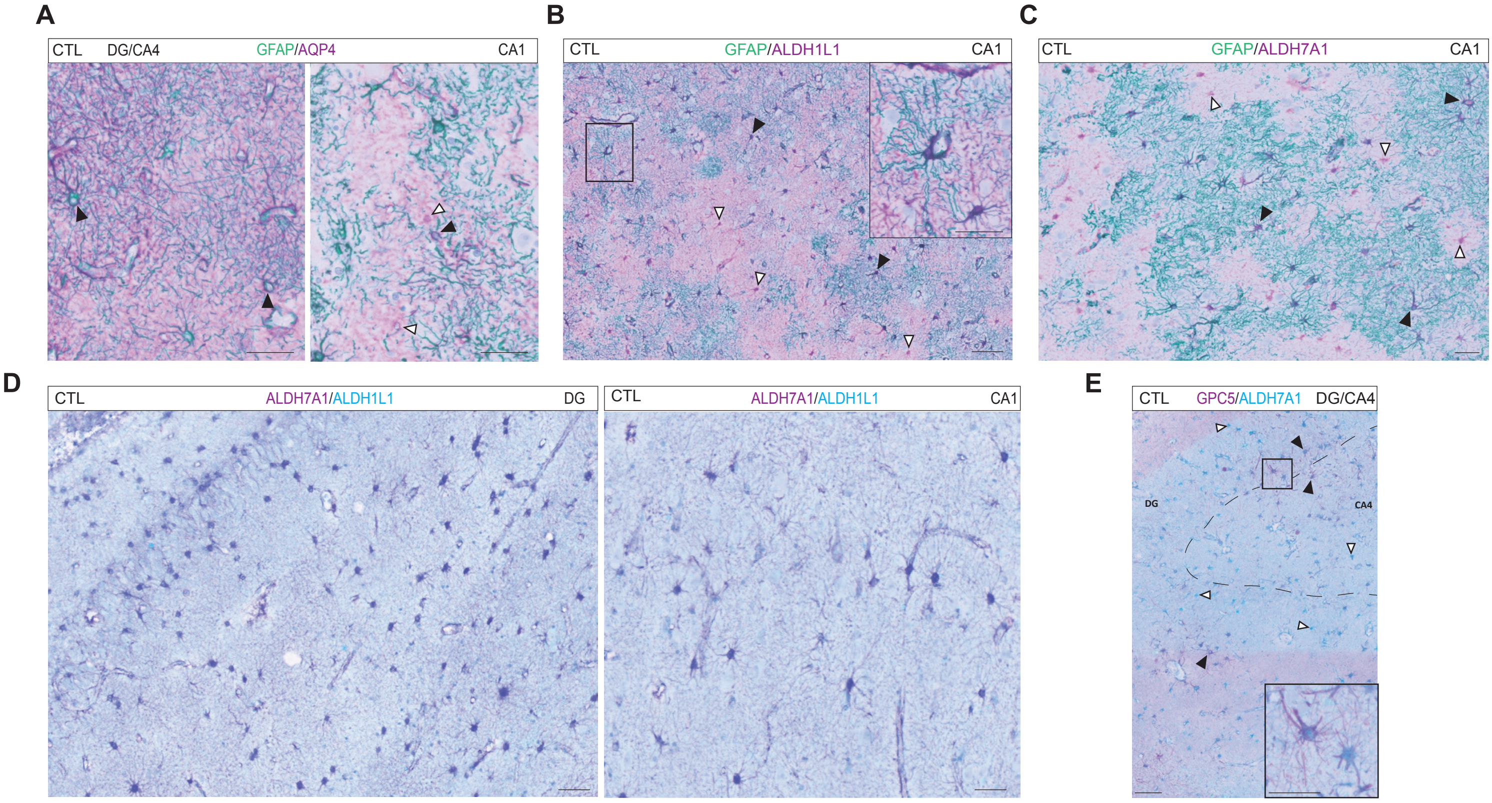


SUPPLEMENTARY FIGURE 3

### Supplementary Figure 4

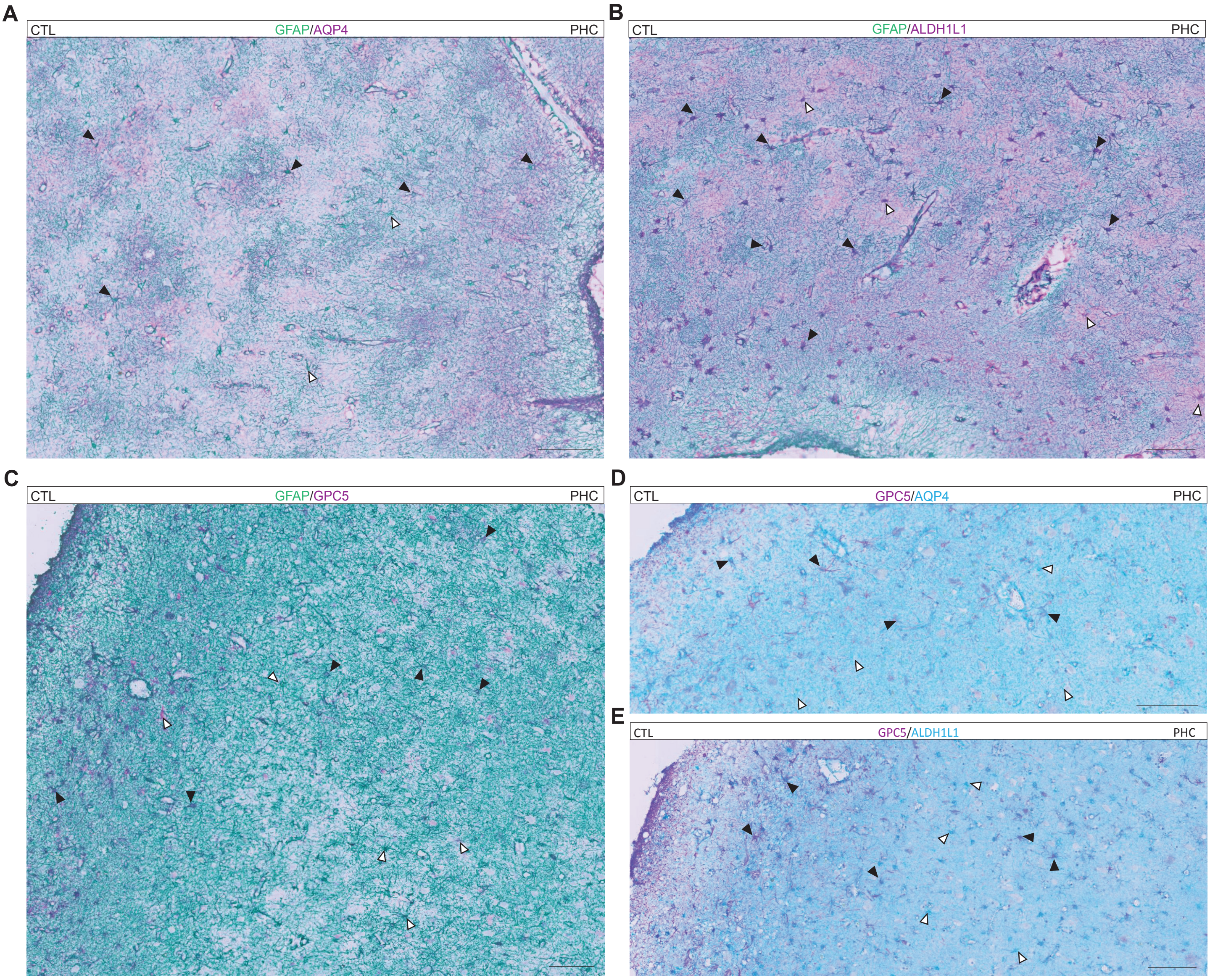


**SUPPLEMENTARY FIGURE 4**

### Supplementary Figure 5

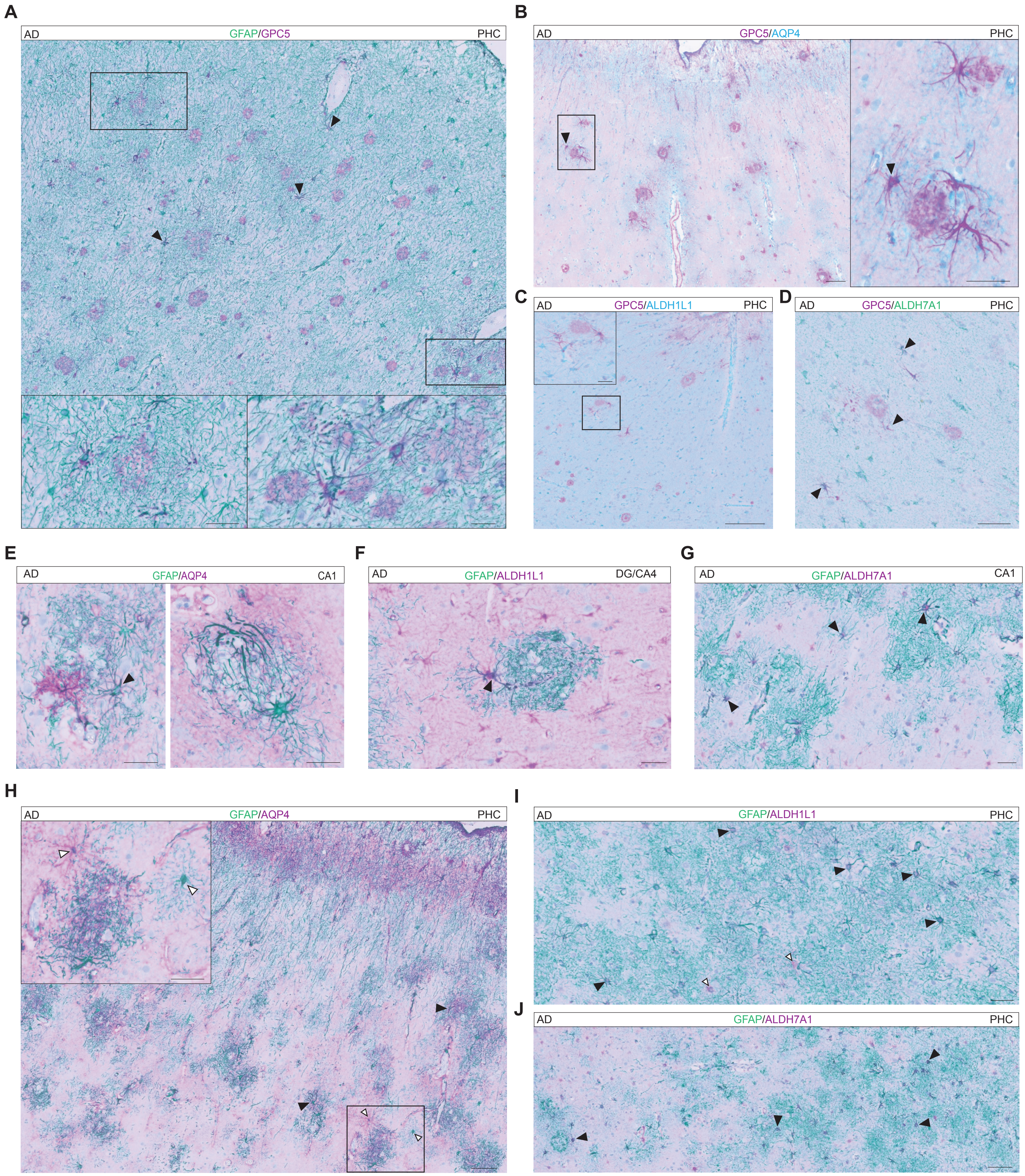


**SUPPLEMENTARY FIGURE 5**

### Supplementary Figure 6

A

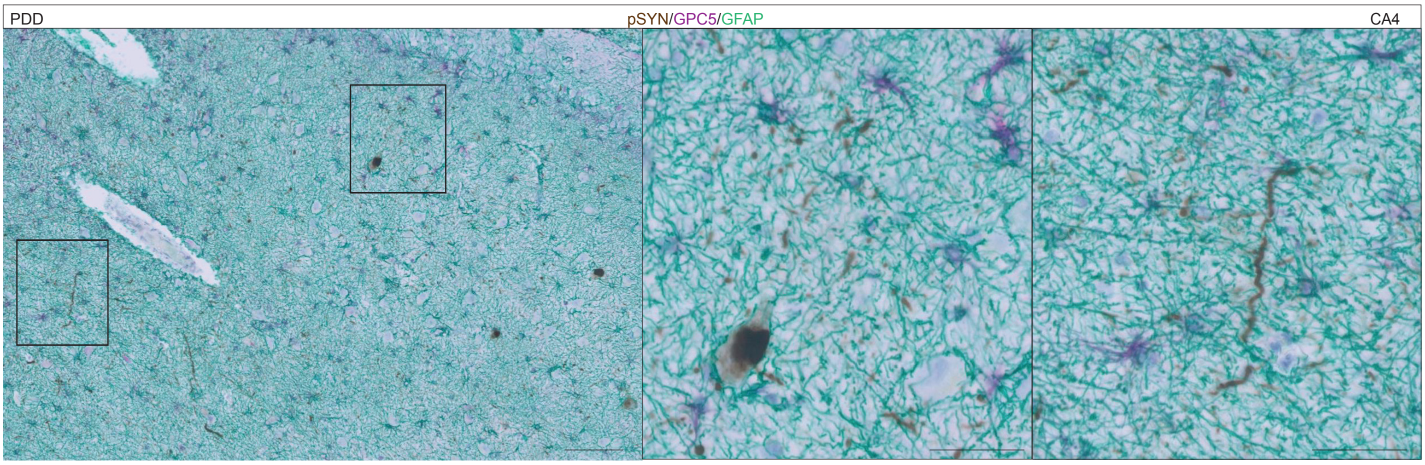


B

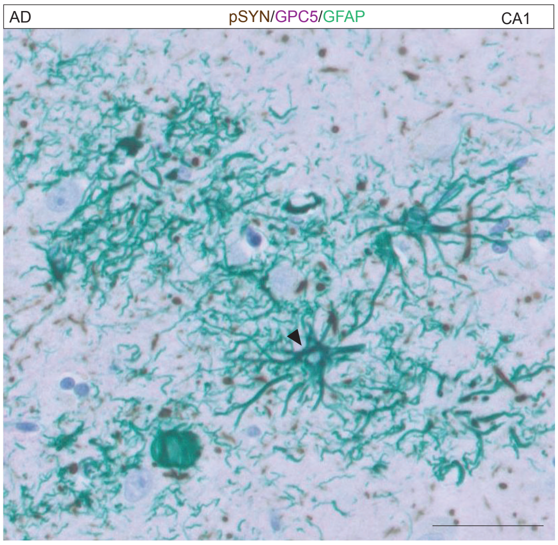


**SUPPLEMENTARY FIGURE 6**
