## Supplementary File 1 for "Glypican-5 delineates spatially distinct astrocyte and neuronal populations in the human hippocampus and is selectively remodelled in Alzheimer’s disease"

Laboratoire national de santé (LNS)

1, Rue Louis Rech

L-3555 Dudelange

**Supplementary Fig. 1: Distribution of GPC5, GFAP, AQP4 and ALDH1L1 in the parahippocampal cortex of healthy aging brain**

**A** In the PHC, GPC5-positive astrocytes (DAB, brown) are mainly found in the glia limitans and the first and second cortical layers (full arrows), while GPC5-positive neurons are sparse (asterisks) (case #6). **B** GFAP-positive astrocytes (DAB, brown) are differentially distributed across the PHC layers. In the deeper layers, GFAP+ astrocytes are organised into non-overlapping domains with many expressing high levels of GFAP ($GFAP_{HIGH}$, full lines) and some expressing low levels ($GFAP_{LOW}$, dashed lines) (case #6). **C** AQP4 (DAB, brown) distribution shows a PHC stratification with an enrichment in the glia limitans, the second and deeper layers and displayed a strong perivascular expression (case #6). **D** ALDH1L1 chromogenic staining (DAB, brown) shows that the first PHC layer is less enriched than the deeper layers (case

#6). Scale bars: **A** low magnification: 100 μm, high magnification: 50 μm; **B-D** low magnification: 200 μm, high magnification: 50 μm.

**Supplementary Fig. 2: ALDH7A1 is a generic marker for hippocampal astrocytes.**

**A** Representative IHC stainings of ALDH7A1 (DAB, brown) in the healthy hippocampus. **a-d** Zoom-in on the different hippocampal subfields: (**Aa**) DG-CA4, (**Ab**) CA3, (**Ac**) CA1 and (**Ad**) subiculum (case #7). **SO**, stratum oriens; **SPy**, stratum pyramidale; **SR**, stratum radiatum. Scale bars: low magnification: 500 μm, high magnification (a-d): 50 μm.

**Supplementary Fig. 3: Astrocytes of the hippocampus display a spectrum of identities.**

**A** Multiplex cIHC for GFAP (green) and AQP4 (purple) revealing a large number of GFAP/AQP4 double-positive astrocytes (dark purple, full arrows), which are found amongst AQP4-positive cells (white arrows) (case #9). **B** Co-staining of GFAP (green) and ALDH1L1 (purple, white arrows) showing a widespread expression of ALDH1L1 across the astrocyte population through the hippocampal layers. In contrast, GFAP labels only specific subgroups of astrocytes that co-express ALDH1L1 (dark purple, full arrows) (case #9). **C** ALDH7A1 (purple, white arrows) is widely expressed by hippocampal astrocytes. GFAP-positive astrocytes (green) also co-express ALDH7A1 (dark purple, full arrows) (case #9). **D** Multiplex cIHC of ALDH7A1 (purple) with ALDH1L1 (teal) showing that hippocampal astrocytes co-express both markers (dark blue) (case #6). **E** GPC5-positive astrocytes are positive for ALDH7A1 (dark blue, full arrows) (case #12). Scale bars: **A, C, D:** 50 μm. **B, E** low magnification: 100 μm, high magnification: 50 μm.

**Supplementary Fig. 4: Astrocytes of the parahippocampal cortex display a spectrum of identities.**

**A** In the healthy PHC, most of GFAP+ (green) astrocytes co-express AQP4 (dark purple) (case #9). **B** They also represent a subpopulation of ALDH1L1+ astrocytes (dark purple) (case #9). **C** The glia limitans is composed of double-positive GFAP+/GPC5+ astrocytes (dark purple) that can also be observed in the first cortical layer. Deeper layers, however, are mainly populated by single GFAP+ astrocytes (green) (case #12). **D** GPC5+ (purple) astrocytes of the PHC also express (**D**) AQP4 and (**E**) ALDH1L1 (dark blue) (case #12). White and black arrows indicate single and double-positive astrocytes, respectively. Scale bars: 100 $\mu$m.

**Supplementary Fig. 5: Hippocampus and PHC astrocytes exhibit homeostatic and disease signatures in AD patients.**

**A-D** The GPC5 co-stainings reveal GPC5+ astrocytes associated to plaque-like structures co-expressing (**A**) GFAP (dark purple), (**B**) AQP4 (dark purple), (**C**) ALDH1L1 (dark blue) and (**D**) ALDH7A1 (dark purple) (cases #28, #29, #33, #34). **E-G** Multiplex cIHC of (**E**) GFAP (green) with AQP4 (purple), (**F**) GFAP (green) with ALDH1L1 (purple) and (**G**) GFAP (green) with ALDH7A1 (purple) in AD samples shows that GFAP astrocytes still co-express AQP4, ALDH1L1 and ALDH7A1 (dark purple, black arrows). AQP4+ and ALDH1L1+ astrocytes are found close to plaques while ALDH7A1+ cells are mainly observed around astrocyte tangle mesh (cases #28, #33, #34). **H-J** Multiplex cIHC of GFAP (green) with (**H**) AQP4 (purple) or (**I**) ALDH1L1 (purple) or (**J**) ALDH7A1 (purple) in the PHC of AD patients highlights the mosaic pattern of expression of these markers (cases #33, #34). White and black arrows indicate single and double-positive astrocytes, respectively. Scale bars: **A-C, H**: low magnification: 100 μm, high magnification: 50 μm; **D-G, I-J**: 50 μm.

**Supplementary Fig. 6: GPC5 is not associated with alpha-synuclein pathology in PDD samples.**

**A-B** In PDD, multiplex cIHC of GPC5 (purple) with GFAP (green) and pSyn (DAB, brown) showed no direct association between GPC5 and Lewy bodies or neurites (**A**), but revealed some GFAP-positive astrocytes bearing intracellular pSyn in CA1 (**B**, black arrows) (cases #42, #51). Scale bars: **A** low magnification: 100 μm, high magnification: 50μm; **B** 50μm.

**Supplementary Table 1: Patient information of the samples used in this study.** Details regarding the samples used in this study and their corresponding neuropathological reports were obtained from brain banks.

| Pathological diagnosis | Case | Sex | Age at death (years) | PMD (hh:mm) | ABC / McKeith score/ Braak Lewy Bodies | Brain Bank |
|---|---|---|---|---|---|---|
| CTL | 1 | F | 83 | 35:45 | A0B0C0 | Douglas Canada Bell Brain Bank |
| | 2 | M | 83 | 16:50 | A0B0C0 | |
| | 3 | M | 72 | 07:29 | A0B0C0 | |
| | 4 | M | 68 | 23:22 | A0B1C0 | |
| | 5 | M | 87 | 04:50 | A2B1C0 | |

| | | | | | | |
|---|---|---|---|---|---|---|
| | 6 | F | 91 | 09:30 | A0B2C0 | The Netherlands Brain Bank |
| | 7 | M | 79 | 06:20 | A1B1C0 | |
| | 8 | F | 102 | 03:55 | A1B2C0 | |
| | 9 | F | 79 | 06:00 | A2B2C2 | |
| | 10 | M | 84 | 05:20 | A1B1C0 | |
| | 11 | F | 86 | 06:25 | A2B1C1 | |
| | 12 | F | 104 | 07:33 | A2B2C1 | |
| | 13 | M | 91 | 06:35 | A1B1C0 | |
| | 14 | F | 92 | 06:00 | A3B2C2 | |
| | 15 | F | 90 | 07:30 | A1B2C0 | |
| AD | 16 | F | 83 | 25:00 | A2B3C2 | Douglas Canada Bell Brain Bank |
| | 17 | M | 91 | 25:00 | A2B3C2 | |
| | 18 | F | 96 | 26:30 | A2B3C2 | |
| | 19 | F | 81 | 23:45 | A2B3C2 | |
| | 20 | M | 77 | 19:25 | A2B3C2 | |
| | 21 | M | 90 | 14:36 | A2B2C2 | |
| | 22 | M | 90 | 26:07 | A3B2C3 | |
| | 23 | F | 87 | 17:00 | A1B2C2 | |
| | 24 | M | 90 | 31:59 | A2B2C3 | |
| | 25 | F | 86 | 39:00 | A3B3C2 | GIE-Neuro-CEB biobank |
| | 26 | M | 85 | 21:00 | A3B1C1 | |
| | 27 | F | 84 | 24:00 | A3B3C2 | |
| | 28 | F | 53 | 06:30 | A3B3C3 | The Netherlands Brain Bank |
| | 29 | F | 70 | 07:25 | A3B3C3 | |

| | | | | | | |
|---|---|---|---|---|---|---|
| | 30 | F | 86 | 03:50 | A3B3C3 | |
| | 31 | M | 38 | 05:45 | A3B3C3 | |
| | 32 | F | 92 | 05:15 | A3B3C3 | |
| | 33 | F | 98 | 05:45 | A2B2C2 | |
| | 34 | M | 64 | 04:58 | A3B3C3 | |
| | 35 | M | 74 | 05:25 | A3B3C3 | |
| | 36 | F | 86 | 06:00 | A3B3C3 | |
| | 37 | M | 57 | 04:55 | A3B3C3 | |
| | 38 | F | 71 | 09:25 | A3B3C3 | |
| | 39 | M | 70 | 04:50 | A3B3C3 | |
| | 40 | M | 75 | 07:45 | A3B3C3 | |
| | 41 | M | 79 | 04:10 | A3B2C1 | |
| PDD | 42 | M | 71 | 04:35 | A1B1C0/LB5 | The Netherlands Brain Bank |
| | 43 | M | 73 | 05:35 | A1B1C0/LB5 | |
| | 44 | F | 83 | 06:05 | A0B1C0/LB5 | |
| | 45 | M | 72 | 04:00 | A0B1C0/LB4 | |
| | 46 | M | 61 | 05:00 | A1B1C0/LB6 | |
| | 47 | F | 71 | 09:05 | A0B1C0/ LB5 | |
| | 48 | M | 85 | 05:15 | A1B2C1/ LB4 | |
| | 49 | F | 73 | 06:10 | A1B1C0/ LB4 | |
| | 50 | F | 88 | 06:05 | A1B1C0/ LB6 | |
| | 51 | F | 69 | 07:05 | A1B1C0/ LB6 | |

**Supplementary Table 2: Antibodies used in IHC chromogenic** (Dako Omnis and Ventana Discovery Ultra).

| **Antibodies** | **Source** | **Identifier** | **Dilution IHC** | **Dilution cIHC** |
|---|---|---|---|---|
| **PRIMARY ANTIBODIES** | | | | |
| Rabbit polyclonal anti-ALDH1L1 | Atlas Antibodies | Cat# HPA050139, RRID: AB_2681031 | 1:2000 | 1:2000 |
| Rabbit polyclonal anti-ALDH7A1 | Sigma-Aldrich | Cat# HPA023296, RRID: AB_1844738 | - | 1:1000 |
| Mouse monoclonal anti-phospho tau (AT8) (Ser202,Thr205) | Thermo Fisher Scientific | Cat# MN1020, RRID: AB_223647 | - | 1:1000 |
| Rabbit polyclonal anti-phospho tau (ps396) | Thermo Fisher Scientific | Cat# 44-752G, RRID: AB_2533745 | - | 1:1000 |
| Mouse monoclonal anti-AQP4 | Atlas Antibodies | Cat# AMAb90537, RRID: AB_2665579 | 1:200 | 1:200 |
| Rabbit monoclonal anti-GFAP | Ventana Medical Systems | Cat# 760-4345 (also 05269784001), RRID: N/A | - | Ready to use* |

| | | | | |
|---|---|---|---|---|
| Rabbit polyclonal anti-GPC5 | Atlas Antibodies | Cat# HPA040152, RRID: AB_2676863 | - | 1:500 |
| Mouse monoclonal anti-phospho synuclein (clone 81A, pSYN) (Ser129) | Millipore | Cat# MABN826 RRID: AB_2904158 | - | 1:500 |
| Rabbit polyclonal anti-VGAT | Sigma-Aldrich | Cat# HPA058859, RRID: AB_2683836 | - | 1:200 |
| Rabbit polyclonal anti-VGLUT1 | Atlas Antibodies | Cat# HPA063679, RRID: AB_2685086 | - | 1:200 |
| Mouse monoclonal anti-β-Amyloid, AA17-24 (4G8) | BioLegend | Cat# 800712, RRID: AB_2734548 | - | 1:2000 |
| Mouse monoclonal anti-NeuN (clone A60) | Sigma-Aldrich | Cat# MAB377, RRID: AB_2298772 | - | 1:200 |
| **SECONDARY ANTIBODIES** | | | | |
| DISCOVERY OmniMap anti-Rb HRP | Roche | Cat# 760-4311, RRID: AB_2811043 | - | Ready to use* |

| | | | | |
|---|---|---|---|---|
| DISCOVERY OmniMap anti-Ms HRP | Roche | Cat# 760-4310, RRID: AB_2885182 | - | Ready to use* |
| DISCOVERY CM DAB kit | Roche | Cat# 760-159, RRID: N/A | - | Ready to use* |
| DISCOVERY Purple Kit | Roche | Cat# 760-229, RRID: N/A | - | Ready to use* |
| DISCOVERY Teal HRP kit | Roche | Cat# 760-247, RRID: N/A | - | Ready to use* |
| Discovery Green HRP kit | Roche | Cat#760-271, RRID: N/A | - | Ready to use* |
| Discovery mRNA Red detection kit | Roche | Cat# 760-234, RRID: N/A | - | Ready to use* |
| **RNAScope ISH PROBES** | | | | |
| RNAscope 2.5 VS Probe – Hs-GPC5 | ACD/Bio-Techne | Cat# 521731, RRID: N/A | - | Ready to use* |
| RNAscope™ 2.5 VS Negative Control Probe-Hs-DapB | ACD/Bio-Techne | Cat# 312039, RRID: N/A | - | Ready to use* |
| RNAscope™ 2.5 VS Positive Control Probe-Hs-PPIB | ACD/Bio-Techne | Cat# 313909, RRID: N/A | - | Ready to use* |

*Vials ready-to-use purchased from Roche Ventana Medical Systems
